# Highly multiplexed community profiling of sub-nanoliter droplet-based co-cultures

**DOI:** 10.1101/2025.01.14.633022

**Authors:** James Y. Tan, Jessica D. Li, Auden Bahr, Xiaoxia Nina Lin

**Affiliations:** Department of Chemical Engineering, University of Michigan, Ann Arbor, MI; Department of Microbiology and Immunology, University of Michigan, Ann Arbor, MI; Gilbert S. Omenn Department of Computational Medicine and Bioinformatics, University of Michigan, Ann Arbor, MI; Department of Biomedical Engineering, University of Michigan, Ann Arbor, MI

## Abstract

A systems approach to microbial community ecology requires high-throughput tools for identifying key microbial interactions. While microfluidic droplets enable the high throughput generation, co-cultivation, and sorting of miniaturized co-cultures in sub-nanoliter, uniform water-in-oil emulsions, analyzing composition of individual droplet co-cultures at comparable throughputs has been challenging, particularly with environmentally isolated or less genetically tractable strains. To address this bottleneck, we present Cocoa-seq (<u>co</u>mbinatorial <u>co</u>-cultivation and <u>a</u>mplicon <u>seq</u>uencing), a droplet-based microfluidic workflow that enables 16S rRNA gene amplicon sequencing for thousands of droplet co-cultures in parallel. In summary, after co-cultivation of microbial co-cultures in agarose droplets, which are then set and recovered as discrete gel beads, Cocoa-seq pairs individual gel beads with barcoded primer beads in new droplets to produce multiplexed amplicon libraries via droplet PCR for sequencing. To benchmark Cocoa-seq, we used a model two-species co-culture and four-member mock communities with three different compositions. The community profiles derived from Cocoa-seq were qualitatively consistent with fluorescence microscopy-based estimates for the two-species co-culture and correlated well with mock community expectations, even for low-abundance representatives. Attempts to incorporate spike-in 16S standards for quantification of absolute abundance were hindered by PCR stochasticity at the single-molecule level, and we provide a simulation-based explanation of this bias and recommend against relying on spike-ins for quantitative inference.

**Importance:** Researchers in microbial ecology are increasingly utilizing sub-nanoliter microfluidic droplets to construct, grow, sort, and analyze miniaturized microbial co-cultures to identify and characterize critical microbial interactions. However, while the generation, incubation, and sorting of droplet co-cultures is scalable, analyzing the composition of these droplet co-cultures in large number is much more difficult. We demonstrate and benchmark a workflow for multiplexing 16S rRNA amplicon libraries derived from thousands of individual droplet co-cultures in a single sequencing run. This addition in the arsenal of growing microfluidic capabilities will support future study of critical interactions within complex microbial systems.

**Study funding:** National Science Foundation (Awards 2120909, 2426415) (awarded to XNL)

## Introduction

The contribution of microbiomes to host-associated (1–3) and environmental health (4–6) is becoming clearer, and ways to predict and manage them are increasingly important. Along with environmental factors, microbe-microbe interactions are hypothesized to be major driving factors of community dynamics, especially at more resolved scales (7). For example, mutualistic cross-feeding networks (8, 9) and microbial antagonism – such as bacteriocin production (10) or Type VI secretion (11) – influence community composition and function. However, identifying and measuring these interactions in complex microbial systems is challenging (12).

There are two major approaches for studying microbial community interactions: top-down, culture-independent ‘omics and bottom-up, reductionist synthetic ecology studies. On one hand, the broad applicability of culture-independent methods has generated large 16S amplicon and ‘omics datasets to infer interactions computationally through statistical correlations (13, 14), genome-scale models (15, 16), or multi-omics (17). While these approaches are scalable, the appropriate empirical data to validate or improve them remains challenging to obtain. On the other hand, synthetic ecology involves interrogating laboratory co-cultures and is capable of generating high-resolution phenotypic and functional data (18). This reductionism has enabled strides in the design of synthetic microbial communities and study of their functions (19–21).

Since the total number of co-culture combinations for a community of *n* members is 2*^n^* - 1, these studies use multiwell plates, microfluidics, or a liquid handling robot, for communities of 6, 19, and 25 members, respectively, to overcome the challenge of combinatorial scalability. However, the major technical barriers to implementing synthetic ecology for a microbial system are the laborious nature and selective bias of laboratory isolation and the throughput-limited manual construction of co-cultures.

The miniaturized nature of microfluidic droplets shows potential to bridge these top-down and bottom-up approaches through the scalable generation and analysis of co-cultures. Microfluidic droplets – uniform, sub-nanoliter water-in-oil emulsions - circumvent traditional co-culture handling by co-encapsulating and co-cultivating random combinations of cells from a sample in high throughput (22–24). Additionally, individual droplet co-cultures can be sorted in high-throughput for functional properties, such as high secretion phenotypes (25, 26), inhibition (27–29), or facilitative interactions (30). However, the current bottleneck is the throughput of droplet co-culture analysis, particularly determining the composition of droplet co-cultures. Prior work has individually selected a low number of droplet co-cultures for manual isolation or sequencing (23, 27–29). Previous droplet-based workflows capable of high-throughput analysis for studying microbial interactions require fluorescent labeled strains (24) or droplet-color coding (20). However, these requirements severely limit the degree of complexity and range of microbes that can be studied due to genetically tractability or laboratory isolation.

To address this bottleneck, we developed Cocoa-seq (<u>co</u>mbinatorial <u>co-</u>culturing and <u>a</u>mplicon <u>seq</u>uencing), which enables high-throughput analysis of droplet co-cultures via droplet-based multiplexing for sequencing. Inspired by droplet-enabled workflows for single cell sequencing (31, 32), single cell transcriptomics (33–36), and 16S amplicon-based spatial mapping (37), we utilize hydrogel beads to enable multi-step processing of droplet co-cultures and massively parallel barcoding. To validate Cocoa-seq, we characterized the quality of droplet-derived amplicon libraries and benchmarked the workflow’s ability to accurately measure community composition with a model synthetic co-culture and various mock communities. We also describe attempts to improve Cocoa-seq by incorporating unique molecule standards for absolute quantification and how PCR stochastic bias at the discrete template level prevents this. Finally, we provide recommendations for applying Cocoa-seq in experimental workflows and technical improvements for further optimization.

## Materials and Methods

### Lithography

SU-8 mastermolds with microfluidic device features were generated with standard SU-8 photolithography procedures in a cleanroom environment. Both PDMS microfluidic devices for the generation of agarose beads and high-throughput sample multiplexing are detailed in Fig S1 and were generated with standard soft lithography methods. Full details are in the Supplementary Information.

### Microbial cultures

For benchmarking, we used six microbial cultures: *E. coli* K12 BW25113 (ΔmetA mCherry), *B. subtilis* 168 (Pveg-GFP), *B. subtilis* 168 (Phyperspank-GFP), *Pseudomonas putida* KT2440, *B. thetaiotaomicron* VPI-5482, and *L. crispatus* ATCC 33820. Each overnight cell culture was washed twice, resuspended in PBS, and quantified by counting with a disposable haemocytomer under phase contrast microscopy. The *E. coli* and *B. subtilis* 168 (Phyperspank-GFP) strains were used together as an obligately-syntrophic, two-species model co-culture similar to the one used in Hsu et al. (24). When co-cultivated in droplets, the *E.* coli/*B.subtilis* co-culture was grown in M9. Three 4-member mock communities were comprised of *E. coli*, *B. subtilis* 168 (Pveg-GFP), *P. putida*, and *B. thetaiotaomicron* in different ratios. Details on media and culture conditions are in the Supplementary Information.

### Encapsulation of cells in agarose beads

Droplet generation was performed in a modified large oven incubator (VWR Scientific 1535) to maintain a temperature between 37-40°C to keep low-melting agarose suspensions fluid (Fig S2). Cell suspensions were mixed with agarose solution to obtain the desired cell concentrations and 1.5% agarose solution. Syringe pumps were used to flow oil (HFE7500 Novec engineering oil with 2% fluorosurfactant) and aqueous suspensions through PFTE tubing into the microfluidic device at rates of 3 µL/min each. Droplet emulsions were collected in 1.5 mL microcentrifuge tubes. For co-cultivation, the emulsions were transferred to a 37°C incubator for 20-24 hours. Agarose in droplets was set by placing the emulsions in ice for 10-20 minutes. To collect agarose beads from droplets, we washed the emulsion with PBS-wash buffer and 20% (v/v) perfluorooctanol in HFE7500 oil, washed with 1% (v/v) Span-80 in hexane, and performed final washes in PBS-wash. Full details are in the Supplementary Information.

### Microscope imaging of droplets

For quantification of *E.* coli/*B.subtilis* droplet co-culture community abundances, fluorescence images of hundreds of droplets were taken after co-cultivation. Images were processed in ImageJ, and custom MATLAB scripts were used to quantify total fluorescence from both strains in individual droplets to estimate relative abundance. Full details are in the Supplementary Information.

### Cell lysis in agarose beads

Agarose beads were washed with 1 mL 10 mM Tris-HCl three times, suspended in lysis buffer (1 mM DTT, 1 mM Tris-HCl pH 8.0, 2.5 mM EDTA, 100 mM NaCl, 0.8% lysozyme), and incubated in a shaker at 37°C overnight. Beads were then washed in 10 mM Tris-HCl pH 8.0, suspended in digestion buffer (30 mM Tris-HCl pH 8.0, 10 mM EDTA, 0.8% Triton X-100 (v/v), 0.5% SDS, 1 ug/µL proteinase K), and incubated for 30 minutes at 50°C. To deactivate proteinase, the beads were washed in 10 mM Tris-HCl pH 8.0, 10 mM EDTA, 0.1% Tween-20 (v/v), 5 mM phenylmethylsulfonyl fluoride (PMSF) (Sigma-Aldrich 93482), washed in TET buffer (10 mM Tris-HCl pH 8.0, 10 mM EDTA, 0.1% Tween-20 (v/v)) five times, and stored at 4°C until use. To check that genomic DNA was immobilized in the microgels after cell lysis, a subset of agarose beads was stained with SYBR Green I and visualized with fluorescence microscopy. Full details are in the Supplementary Information.

### Barcoded hydrogel bead processing

Barcode hydrogel beads were purchased as a unit of 1 million suspended in TET buffer (RAN Biotechnology, CustomSeqReady-1M, InDrop variation, ca. 147k). 16S 515f primers were extended onto the barcode beads by complementary extension oligos based off a protocol from Zilionis et al. (34). Beads were washed extensively with multiple buffers to stop extension and remove extension oligos. Any unelongated oligonucleotides on the bead were removed by ExoI clean-up (Thermo Fisher Scientific, EN0581). Final beads are washed and stored at 4°C. After the extension protocol, barcode beads were checked to ensure proper 515f extension through the hybridization of complementary fluorescent oligo probes and visualization with fluorescence microscopy. Full details are in the Supplementary Information.

### 16S multiplexing

515f-extended barcode beads, gDNA-agarose gels at a 50% (v/v) suspension, and 2x PCR mastermix solution containing 816r reverse primer, SuperFi II buffer, dNTPs, Platinum SuperFi II DNA polymerase, Pluronic F-68, and bovine serum albumin (BSA) were flowed into the microfluidic device with syringe pumps at rates of 0.45-0.5 µL/min, 0.8-1 µL/min, and 3 µL/min, respectively. Oil (Droplet Generation Oil for EvaGreen, Biorad) was flowed at 5 µL/min. Droplets of around 150 µm diameter were generated. For each sample, 20-40 µL of droplets and 20 µL of fresh Biorad oil were added into PCR tubes and exposed to 365-nm UV-light for 10 minutes on ice. 50 µL of mineral oil was placed on top of the droplets to prevent evaporation, and droplets were thermocycled (95 °C for 2 min; 98 °C for 30 sec; and 30 cycles of 98 °C for 10 seconds, 60 °C for 10 seconds, and 72 °C for 30 sec) with a 2 °C/s ramp rate between all steps with no heated lid. Droplets underwent clean-up, involving droplet breaking with 20% perfluorooctanol in oil, primer degradation with ExoI, and Ampure bead purification. Full details for droplet multiplexing are in the Supplementary Information.

### Unique molecular identifier (UMI) benchmarking

To quantify the amplification of individual 16S template molecules, we designed an oligonucleotide composed of a modified 16S V4 sequence from *Thermus thermophilus* ATCC 33923 with seven nucleotides directly after the forward primer region replaced with seven degenerate nucleotides for use as unique molecular identifiers. The 16S standard was purchased from GenScript as a ssDNA stock, resuspended in molecular-grade water, and quantified with an in-house digital droplet polymerase chain reaction (ddPCR) to determine the concentration of functional 16S standard molecules. The stock was diluted and co-flowed into droplets in the droplet barcoding device and workflow for a desired initial λ of 50 molecules/droplet. Full details for oligonucleotide design and in-house ddPCR protocol are in the Supplementary Information.

### Illumina library preparation and sequencing

Illumina adaptors and indices were attached to each multiplexed library via PCR with NEBNext Q5 HotStart HiFi polymerase. The thermocycler program was as follows: 98 °C for 30 seconds; 5-7 cycles of 98 °C for 10 seconds, 68 °C for 20 seconds, and 65 °C for 30 seconds; 65 °C for 2 minutes; and a 12 °C hold. PCR product was verified with gel electrophoresis and purified with Ampure XP bead clean-up. Libraries were sent to the University of Michigan Advanced Genomics Core for quality control with Agilent TapeStation and sequencing with the Illumina NextSeq 2000 with P1 300 cycle chemistry. Details for library preparation PCR and sequencing details are in the Supplementary Information.

### Bioinformatic processing

A custom demultiplexing Python script (run on v.3.10.4) based on a previous study (34) was modified and used to demultiplex reads and trim barcodes from the forward reads. Because of the large number of barcode families (groups of reads with the same barcode) with a very small number of total reads, we used the expected number of barcodes (Note S1) to filter on the barcode families with the highest number of reads. Using vsearch (v.2.13.7-linux-x86-64), operational taxonomic units (OTUs) were constructed for reads within a 95% similarity threshold to expected 16S v4 sequences for known isolates with the usearch global command (38). To run sample rarefaction and beta-diversity analyses, we used mothur (v.1.44.1) (39). To compare sequencing reads to cell abundances, we normalized abundance by 16S copy number for each strain, retrieved from rrnDB (40). Sequence read processing scripts were run on the Great Lakes HPC Cluster (University of Michigan).

### Stochastic amplification simulation

To simulate the effect of stochastic bias for a low number of initial molecules, we programmed a simple Python script with a model in which individual template molecules were represented uniquely as different characters in an array. The probability of success of amplification in one round of PCR was represented by the Bernoulli distribution in which success of a molecule is given by *η_amp_* (amplification efficiency) and failure is given by 1-*η_amp_* If a molecule was replicated, its unique character string was added to the array. This was propagated for a designated number of cycles for each molecule. At the end, to simulate sequencing, we subsampled the number of molecules to 1000 molecules and the final abundances of molecules were compared to the initial.

## Results

### Workflow for droplet-enabled high-throughput co-culturing and 16S multiplexing

The Cocoa-seq workflow has two parts: (1) generating and processing co-cultures (Fig. 1a) and (2) barcoding them for highly multiplexed sequencing libraries (Fig. 1c). In the first half of the workflow, Cocoa-seq enables multi-step processing of individual droplet co-cultures by co-encapsulating and co-cultivating them in liquid agarose droplets, setting and recovering them as agarose beads, and lysing the immobilized cells within to expose genomic DNA (gDNA) (Fig. 1a). More specifically, during co-cultivation, initial cells are suspended in growth media with molten low melt agarose and randomly encapsulated following Poisson statistics. The average number of encapsulated cells per droplet (λ) is determined by the concentration of the cellular suspension and the size of the droplet generated (41), which is set by flow rates and microfluidic channel dimensions. Generated droplet co-cultures are co-cultivated under appropriate conditions. After co-cultivation, co-cultures are immobilized in hydrogel beads by lowering the temperature to set agarose. The droplet emulsion is then broken, and co-culture beads are washed to remove residual oil and resuspended in an aqueous suspension. The co-culture beads are subject to enzymatic and chemical cellular lysis, which leaves high molecular weight gDNA in the agarose matrix (Fig. 1b). Our simple cell lysis protocol was effective for the majority of the strains we tested (Fig. S3). The second half of the workflow pairs these gDNA-exposed co-culture beads with barcoded hydrogel beads (BHBs) in another droplet co-encapsulation step, which enables the barcoding of 16S bacterial marker genes via droplet PCR (Fig. 1c).

**Fig. 1.**
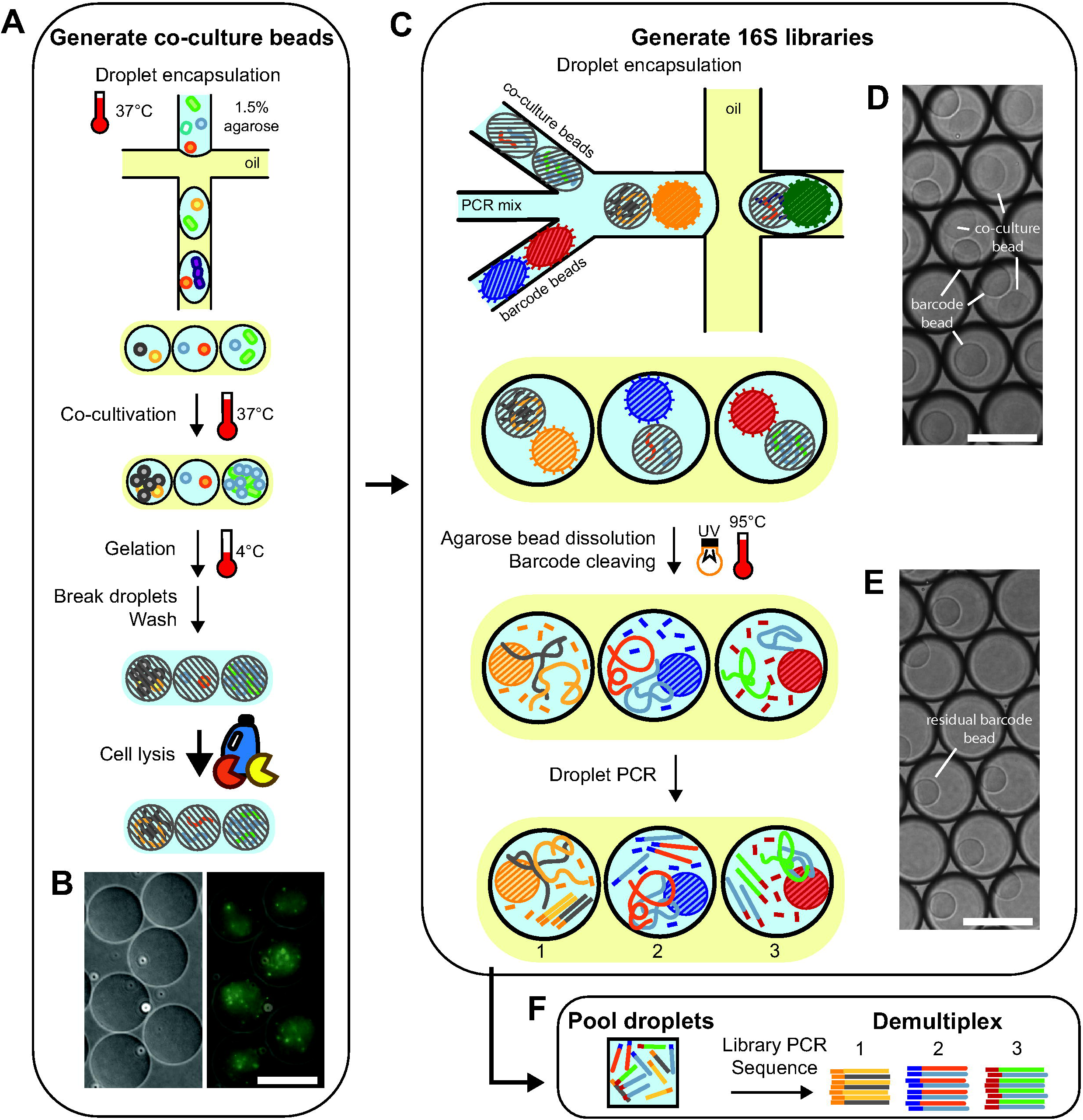
Experimental overview of Cocoa-seq for highly multiplexed 16S amplicon sequencing of droplet co-cultures. (A) Schematic for generation of co-culture agarose beads which hold cells and gDNA of co-cultures generated from random droplet co-encapsulation. (B) Microscope images of agarose beads with gDNA from lysed droplet co-culture cells generated at the end of (A) (left is bright field image, right is SYBR stain overlay). Scale bar is 80 µm. (C) Schematic for multiplexed 16S library generation using co-culture beads from (A) and barcoded hydrogel beads (BHBs) functionalized with bead-unique barcode oligonucleotides. (D) Droplets generated from the co-encapsulation of co-culture beads and BHBs. Polyacrylamide beads are darker in outline and slightly smaller than agarose beads. Scale bar is 150 µm. (E) Droplets after agarose bead dissolution and droplet PCR. The agarose beads have melted while polyacrylamide beads remain. Scale bar is 150 µm. Droplets are not the same subset as those in (D). (F) After pooling droplets and sequencing, bioinformatic demultiplexing of barcodes from BHBs enables identification of reads from each individual droplet.

These BHBs are polyacrylamide hydrogel beads functionalized with oligonucleotides, which are 16S V4 primers with bead-specific unique barcode overhangs in the droplet PCR. To pair individual co-culture beads and BHBs with PCR reagents in droplets, we designed a co-flow microfluidic device which accommodates close-packed flows of two types of hydrogel beads (Note S2) and is demonstrated in the Supplementary Video. While only ∼37% of droplets have both bead types due to microfluidic constraints, ∼82% of co-culture beads are correctly paired (Fig. 1d, Fig. S4), meaning a large majority of the co-cultures are barcoded. In droplets, barcoded oligos are photocleaved from hydrogel beads by UV photoexposure, and agarose beads are melted to release gDNA. Droplet PCR then generates droplet-specific barcoded 16S amplicons. The droplets remain thermostable (Fig. 1e) until intentionally broken in a final bulk limited-cycle PCR to attach Illumina adaptors and indices to the multiplexed amplicons, products are confirmed by size (Fig. S5), the final library is sequenced, and the barcoded 16S amplicon reads are demultiplexed (Fig. 1f).

The BHBs which enable library multiplexing for Cocoa-seq adapts designs from a previous single-cell workflow (34) and requires modification when purchased from vendors (Fig. 2). The oligonucleotides functionalized onto the BHBs have three major components: a photocleavable spacer, a dual-index barcode region, and an extension adaptor. The photocleavable spacer is sensitive to a specific UV-wavelength; when cleaved, it detaches the oligo from the polyacrylamide bead. The dual-index barcode scheme utilizes split-and-pool combinatorial synthesis (34), which enables rapid generation of hundreds of thousands of barcodes. Cocoa-seq utilizes the extension adaptor to incorporate 515f 16S V4 forward primers onto BHB oligos by isothermal polymerase, and the attachment of the 515f primer sequence is confirmed and quantified with probes (Fig. S6). All three components allow these oligos to act as barcoded forward primers for bacterial 16S rRNA marker gene PCR along with the appropriate reverse primer supplied during droplet PCR. The exact oligonucleotide sequence and molecular workflow for multiplexing with BHBs are specified in Fig. S7. BHBs can also be generated in-house according to relevant protocols (34) with appropriate modifications, such as replacing polyT regions originally intended for single cell eukaryotic RNA sequencing.

**Fig. 2.**
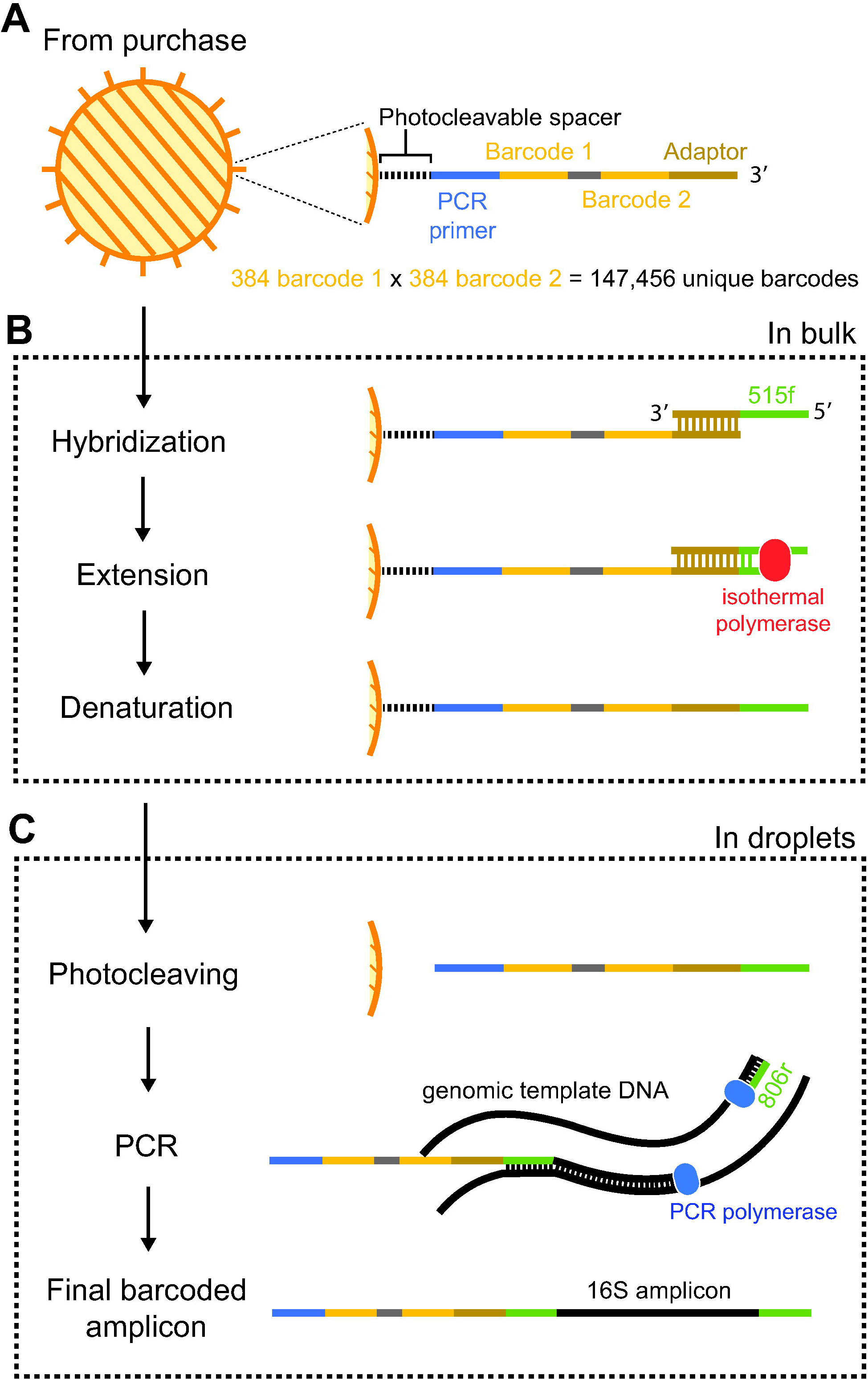
Preparation of barcoded hydrogel beads (BHBs) functionalized with barcode oligonucleotides for Cocoa-seq. (A) Purchased BHBs are functionalized with oligonucleotides with unique bead-specific barcodes through combinatorial dual indices. (B) After purchase, to prepare the beads for Cocoa-seq, oligonucleotides on BHBs are extended in bulk with the 515f primer for amplifying the 16S bacterial universal marker gene. (C) Full oligonucleotides, when cleaved off beads in droplets with a specific wavelength of UV light, act as primers during droplet PCR, resulting in barcoded 16S marker gene amplicons.

### Benchmarking shows barcode filtering is required to remove sequence noise from droplet libraries

To assess the quality of the sequencing libraries generated by Cocoa-seq, we benchmarked with a model bi-culture and a set of synthetic mock communities. To demonstrate Cocoa-seq’s ability to incorporate co-cultivation, we utilized an obligately syntrophic two-species co-culture, comprising of an engineered *E. coli* methionine auxotroph (mCherry-labeled) and a native *B. subtilis* tryptophan auxotroph (GFP-labeled). To ensure the presence of both strains in each droplet, we generated droplets with an initial λ of 5 cells for both species and co-cultivated them overnight, resulting in growth of both (Fig. 3a). Additionally, to demonstrate Cocoa-seq’s ability to accurately represent co-cultures across wider compositional space, we created 3 mock communities comprised of *E. coli*, *B. subtilis*, *P. putida*, and *B. thetaiotaomicron* in different rank abundances and generated a droplet library of random droplet communities for each by co-encapsulation of mixed cellular suspensions with a λ of 100 cells/droplet. The mock community-derived libraries will be referred to as: the “even” (comprised of a 1:1:1:1 λ ratio of each species), the “Ec-biased” (comprised of mostly *E. coli* and the other members, respectively, in decreasing rank abundance), and the “Bt-biased” (with opposite rank composition of the “Ec-biased”) (Fig. 3b).

**Fig. 3.**
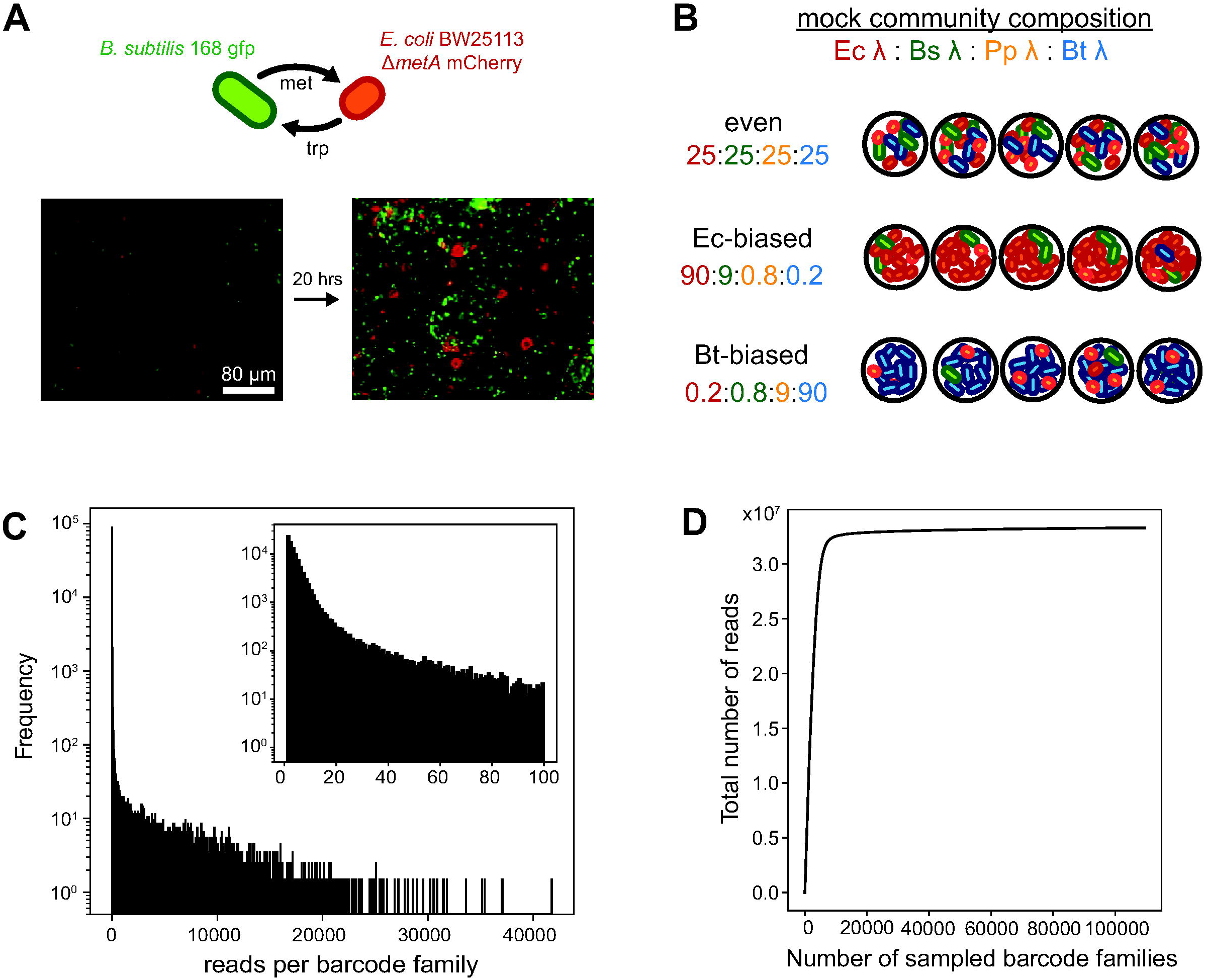
Bi-culture and mock community benchmarking libraries used to validate Cocoa-seq and distribution of barcode family (groups of reads sharing the same barcode) sizes from the bi-culture droplet library. (A) (Top) The model bi-culture is comprised of two cross-feeding species: *B. subtilis* 168, which is a GFP-labeled (green) native auxotroph for tryptophan, and *E. coli* BW25113, which is an mCherry-labeled (red) *metA* knockout mutant. (Bottom) Fluorescence microscopy with brightfield overlays before and after co-cultivation for 20 hours. Initial λ for each species was 5 cells/droplet. Imaged droplets are not the same between timepoints. (B) Three synthetic mock communities were comprised of four members (*E. coli*, *B. subtilis*, *P. putida*, and *B. thetaiotaomicron*) present in different rank abundances with the λ (average cells/droplet) specified in the labelled ratios. Lower rank abundance members occur less than once per droplet. (C) Histogram of the number of reads per barcode family from the bi-culture droplet library with a bin size of 10. The subset histogram shows a more resolved view (bin size of 1) of the smallest barcode families. (D) Rarefaction curve of the cumulative number of reads contributed from sampling barcoding families.

We sequenced these benchmarking libraries generated with Cocoa-seq and assessed library quality through bioinformatic processing (Table 1). After initial Illumina index demultiplexing (Table S1), individual libraries were demultiplexed into individual barcode families (groups of reads with the same barcode), and sequences associated with ambiguous or low-quality barcodes were filtered out. This resulted in 81-83% of total reads with meaningful and differentiable barcodes. There were more barcode families than expected based on the number of droplets processed during droplet barcoding PCR, echoing results from other droplet-based method utilizing a similar microfluidic workflow (37, 42). For example, for the biculture library, which was sequenced the deepest, we observed 109 459 total barcode families, which is much higher than expected around 1 500. The histogram of reads per barcode family in the bi-culture library shows most barcode families contain very few amplicons (Fig. 3c), and this is also observed in the other benchmarking libraries (Fig. S8). A cutoff for the number of reads per barcode family to be considered a genuine barcoding event was set conservatively to match the expected number of co-cultures barcoded (Note S2), keeping anywhere between 50%-80% of the filtered reads. We suspect that “noisy” reads below the barcode family threshold were generated from failed co-encapsulation events (e.g. droplets with only barcode beads), combinatorial barcode index hopping, barcode oligo synthesis error, or amplification error. We do not suspect substantial contamination since a majority of these reads (between 91.3 - 96.4%) were 95% identical to reference sequences. Overall, the total retention rate of reads through bioinformatic processing was between 40% and 65%. Additionally, the sequencing depth of our libraries was intensive enough to capture the available droplet library, as demonstrated by a rarefaction curve of total reads from barcode families for the biculture library (Fig. 3d) as well as the other libraries (Fig. S8).

**Table 1.** Retention of reads through bioinformatic processing for each benchmark library. We removed reads that did not match expected barcode sequences, belonged to barcode families with too few total reads, and did not match expected 16S V4 sequences within 95% identity. The minimum reads/barcode threshold was set based on the number of expected barcodes from the volume of droplets thermocycled during Cocoa-seq (Note S1). we estimated 1500 communities derived from each mock community and 2500 for the bi-culture and made sure to set the threshold to keep less than the estimated number.

| Sample | Barcoding demultiplexing |  | Removing spurious barcode families |  |  |  | 16S classification | Net |
| --- | --- | --- | --- | --- | --- | --- | --- | --- |
|  | Total reads | Reads kept | Total barcodes | Minimum reads/barcode | Barcodes kept | Reads kept | Reads matching | % reads kept |
| even mock | 4 483 938 | 3 709 660<br>(82.7%) | 30 278 | 1 000 | 1 294 | 1 870 277<br>(50.4%) | 1 807 622<br>(96.6%) | 40.3% |
| Bt-biased mock | 17 271 208 | 14 212 426<br>(82.3%) | 59 014 | 4 000 | 1 178 | 11 450 686<br>(80.6%) | 11 118 631<br>(97.1%) | 64.4% |
| Ec-biased mock | 21 314 159 | 17 508 079<br>(82.1%) | 69 669 | 4 500 | 1 485 | 8 915 003<br>(50.9%) | 8 314 130<br>(93.3%) | 39.0% |
| bi-culture | 40 860 298 | 33 296 045<br>(81.5%) | 109 459 | 5 000 | 2 464 | 24 479 988<br>(73.5%) | 22 683 747<br>(92.7%) | 55.5% |

### Cocoa-seq shows qualitative agreement with fluorescence microscopy-based measurements in determining droplet bi-culture composition

We used the model bi-culture to compare community composition distributions between Cocoa-seq and fluorescence microscopy. While fluorescence microscopy is semi-quantitative due to light diffraction at the droplet-oil interface, cell-to-cell fluorescence variation, microscopic plane focusing on a specific two-dimensional cross-section, and cell clumping, we accounted for the first two limitations by normalizing the fluorescence per cell with image processing (Fig. 4a). From this we calculated total fluorescence for both strains to estimate relative abundance across hundreds of droplets. The abundance distributions from both methods are qualitatively comparable, showing the final community state is heavily skewed towards *B. subtilis.* As expected, there is a wide range of final community outcomes (Fig. 4b), which is also observed in other droplet co-culture studies (24, 43), most likely due to stochastic variability in cell-to-cell viability and phenotype. However, the bias towards *B. subtilis* is more pronounced in the sequencing dataset compared to that from fluorescence-based measurements. Given the limitations of fluorescence measurements, we suspect that fluorescence estimates are most likely underestimating *B. subtilis* abundance. The extensive growth and clumping of *B. subtilis*, visible as large green aggregates, were unlikely to have been accurately quantified using two-dimensional cross-section microscopy images. Additionally, 16S PCR bias is reported to introduce a significant degree of systemic sequencing bias, potentially favoring abundant taxa (44–47) and contributing to an overinflated abundance of *B. subtilis*.

**Fig. 4.**
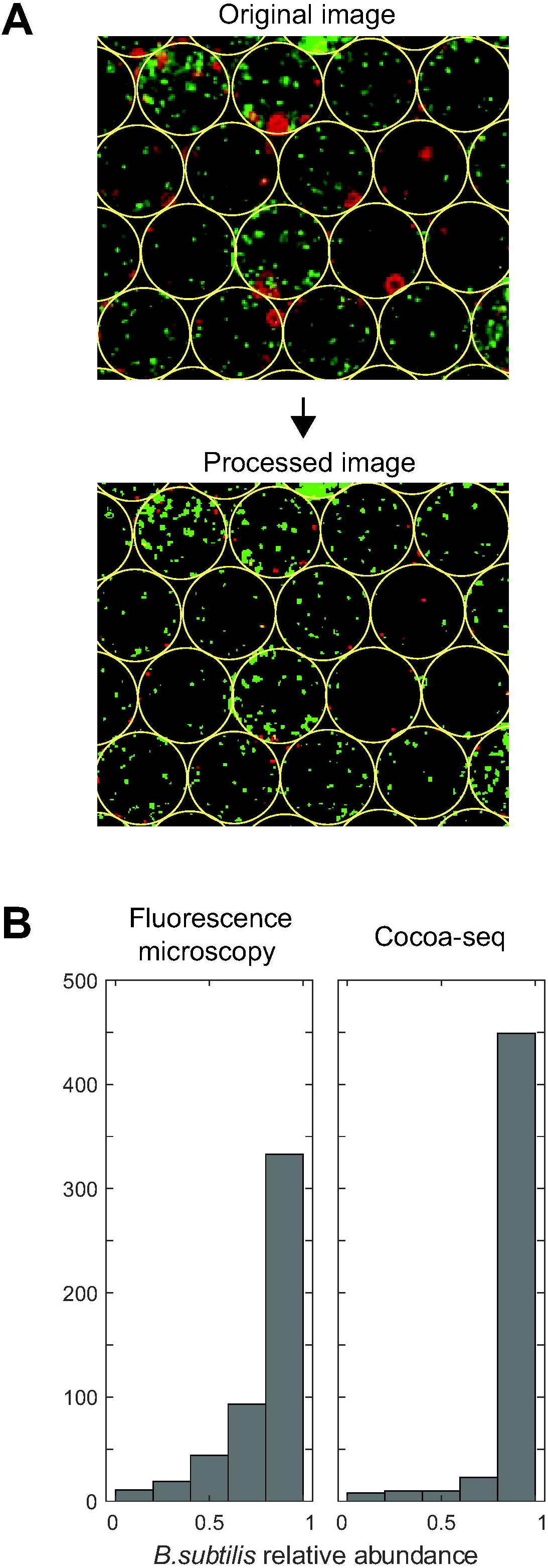
Community composition distributions from fluorescence microscopy and from Cocoa-seq for droplet co-cultures of the syntrophic bi-culture qualitatively agree. (A) Example processing of the fluorescence image from Fig. 3A to detect droplets and normalize fluorescence signal per cell. Droplets (outlined in yellow) are ∼80 µm in diameter. (B) Histograms of *B. subtilis* relative abundance in individual droplets as measured by total fluorescence compared to Cocoa-seq. Each histogram represents a different set of 500 individual droplets from the same droplet pool for comparison.

### Cocoa-seq replicates expected beta-diversity patterns of mock communities libraries

We sought to more rigorously evaluate Cocoa-seq’s ability to measure community composition against more concrete and easily established standards with the previously-described 4-member synthetic mock community libraries. Because there is no cultivation, droplet libraries’ community composition distributions can be simulated from the initial mock community composition before droplet encapsulation. Specifically, we simulated droplet co-encapsulation and sequencing *in silico* based on Poisson statistics and strain-specific 16S rRNA copy numbers. We first compared the distribution of droplet library community compositions from *in silico* expectations to those obtained from Cocoa-seq with principal component analysis (PCA) (Fig. 5a). The communities cluster by droplet library, as expected, but have a higher abundance of *B. subtilis* and a larger spread than in silico distributions, which can be partially explained by cell clumping during droplet encapsulation (Fig. S9a) and the large *B. subtilis* cells used for mock communities, contributing to overrepresentation (Fig. S9b). To determine if Cocoa-seq quantitatively preserves community composition, we determined both the average relative abundance and prevalence (i.e., occurrence rate) across droplet libraries for each member and compared to *in silico* expectations. Both average relative abundance and prevalence largely agree between observation and expectations across high- and low-rank members, showing high linear correlation (R^2^ = 0.854 and 0.946, respectively) (Fig. 5b; Tables S2, S3). This result demonstrates Cocoa-seq meaningfully reports community compositions given the appropriate sequencing depth.

**Fig. 5.**
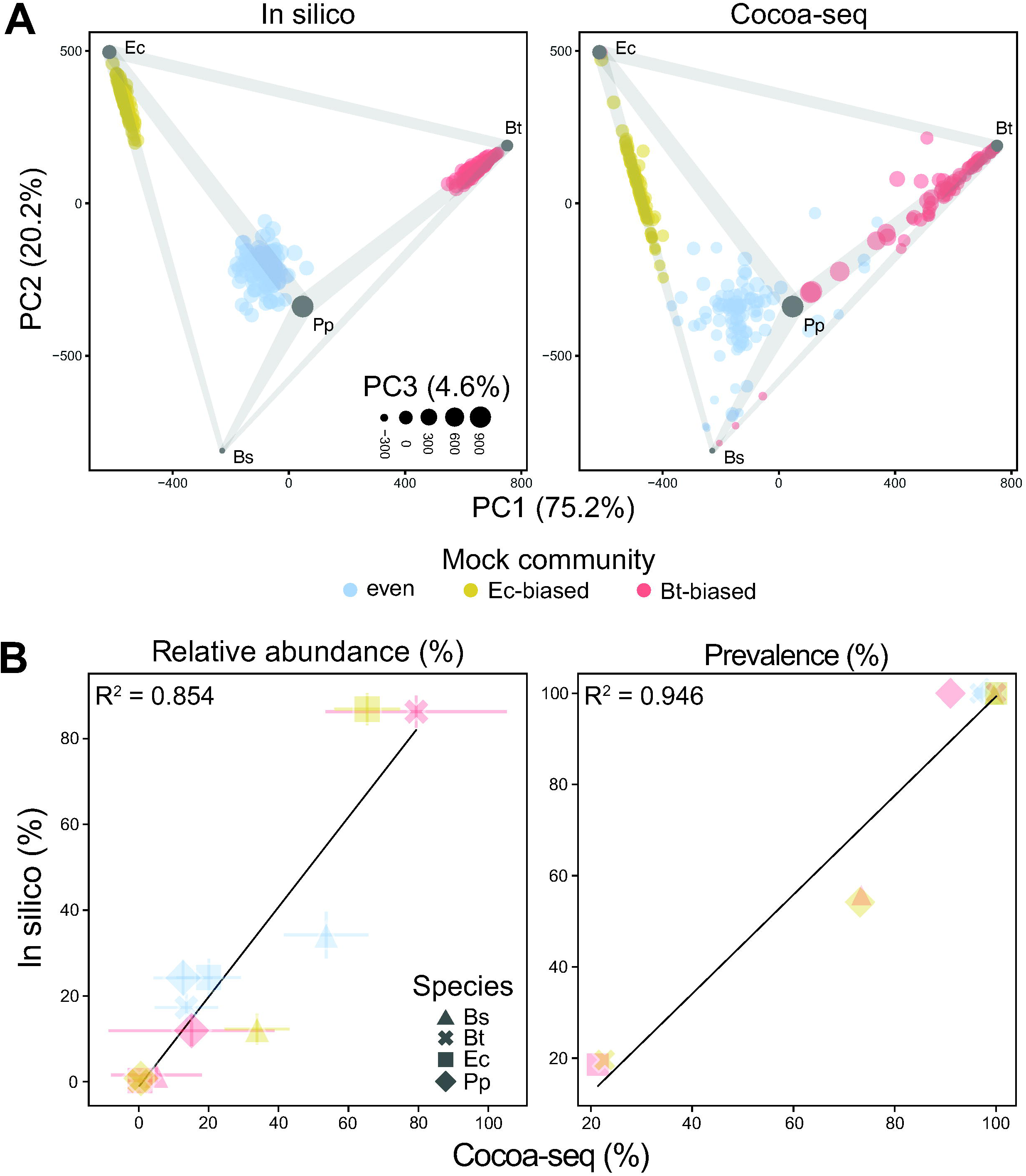
Community composition distributions from Cocoa-seq for droplet libraries agree with expectations. (A) Principal component analysis (PCA) of community compositions in individual droplets derived from a mock community, as expected from simulations based on Poisson statistics and observations with Cocoa-seq (100 instances for each sample). The tetrahedron in 3D (with the third principal component coming out of the page) represents all possible compositions of the four-species ecosystem, with each vertex corresponding to a monoculture (gray circle). Point colors designate the mock community of origin. (B) Linear regression comparing metrics across droplet libraries between Cocoa-seq and the *in silico* expectations. Metrics include the average relative abundance and the prevalence of each member. Since average relative abundance is averaged over all droplets, error bars signify the standard deviation. The color follows the same mock community as in (A) with the marker shapes representing different members. Numerical values for reference are included in Table S2 and Table S3.

### PCR introduces significant bias at single molecule resolution, preventing the use of spike-in molecular standards for absolute quantification

We hypothesized that we could introduce spike-in 16S standards during the barcoding step of Cocoa-seq to infer absolute abundances of droplet communities, enabling the differentiation of total biomass between droplet co-cultures (48) (Fig. S10). We proposed including 16S V4 oligonucleotides with 7-mer random unique molecular identifiers (UMI) (49) (for a total of 4^7^ possible unique molecules) at a λ of 50 molecules/droplet during barcoding PCR and subsequent sequencing. The exact number of spike-in molecules for each droplet would be quantified for each droplet using the total observed UMIs and used to back calculate the absolute abundances of other members. However, when we quantified the amplification of each of these individual spike-in molecules in droplets after Cocoa-seq, there was a very high disparity between discrete template counts despite each template being represented exactly once during PCR (Fig. S11a). Due to this significant variation, determining the total number of spike-in molecules per droplet is difficult due to the inability to determine a reasonable threshold to differentiate signal from noise.

We hypothesized this was a result of stochastic bias at the single molecule level (50, 51). To test this, we simulated PCR with a probabilistic model in which the amplification of an individual template molecule during a PCR cycle was successful based on a binary probability distribution dependent on an arbitrarily chosen amplification efficiency parameter. Iteration of this stochastic model with individual templates over multiple cycles produced a distribution of amplification for individual initial templates (Fig. S11b), which we observed in our data. However, it did not generate a distribution with most templates poorly amplified, as observed in our experiments. There is also stochasticity introduced by subsampling during sequencing and polymerase amplification error that we did not consider in this simple model (52). We explored if the stochastic bias affects the relative abundance profiling of communities by running this model on *in silico* mock communities (Fig. S11c). This stochastic bias is low with higher initial template amounts, but the effect increases as initial template number decreases. While not definitive, they suggest that PCR can introduce significant stochastic bias when amplifying a small number of discrete templates.

## Discussion

Cocoa-seq represents a significant addition in the systems toolbox by enabling the community profiling of thousands of individual droplet co-cultures. Its capacity to be used directly on co-cultivated droplet co-cultures is demonstrated with the model biculture synthetic co-culture, and the quantitative accuracy has been benchmarked against mock communities. We also attempted to enhance Cocoa-seq by introducing spike-in molecule standards for absolute quantification. However, the inherent bias introduced by PCR for single discrete template amplification prevented its use in normalizing read abundance for absolute quantification, which remains a significant technical challenge. Nevertheless, as it is currently presented, Cocoa-seq in conjunction with other droplet-enabled capabilities and experimental designs, such as those discussed in the Introduction, could enable the future study of numerous microbial communities, such as sorting of sub-communities for high biomass to identify mutualistic interactions or for low reporter-strain signal to identify inhibitory interactions.

For some microbial systems, Cocoa-seq may need to be adapted. First, while our lysis protocols were effective for our mock communities, as with all lysis protocols, they must be appropriate for the biological system. The inability to utilize mechanical bead beating to break tough cell wall structures limits universality, but we suspect many other chemical and enzymatic methods can be tested (53). Second, while agarose is a common solid medium gelling agent and works for most systems, it may be problematic in specific situations. For example, in marine systems, agarose is utilized as a natural carbon source (54). In addition, with the current workflow, cultivation temperatures need to be above 37°C to prevent the low melt agarose from setting and constraining cells during cultivation, which matters specifically if cells require physical contact for interaction (55). Environmental microbes also exhibit a very wide range of optimal growth temperatures, and the current inability to accommodate this is another limitation. Ultra-low melt agarose is a feasible alternative since it can remain liquid at room temperature. Alternatively, polyacrylamide could replace agarose, but its monomer toxicity is understudied (56) and it may need to be added to droplet co-cultures after cultivation by microfluidic picoinjection (57). More recently, commercially available semi-permeable capsules enable multi-step processing of microbial cultures (58), and adapting Cocoa-seq with these micro-compartments may enable more universal workflows.

While we believe the highest-impact applications of Cocoa-seq would be to study complex natural microbial communities, Cocoa-seq can also aid the construction and investigation of synthetic communities. Its throughput can accommodate a higher number of strains compared, even to robotics-based approaches using multi-well plates. Additionally, while approaches to high-throughput community design currently rely on linear, objective-based progressions through the community design space (21, 59), the stochastic nature of Cocoa-seq facilitates a broader search of the community design space. Results from Cocoa-seq can inform more precise traditional experimental design by providing more diverse initial points in the design space, which may increase the speed of optimization and prevent trapping in local optima.

Finally, we want to acknowledge that a major challenge for widespread utilization of microfluidic workflows is the need for technical knowhow. To help mitigate the issue, in addition to the Supplementary Information, which details more specifics, we have posted a protocols.io live document with more nuanced experimental specifications and opportunity for inquiry. We also recommend those who are interested in applying Cocoa-seq to leverage commercially available droplet platforms and/or consult groups that utilize microfluidics at their own or neighboring institutions.

## Acknowledgements

The authors would like to acknowledge helpful conversations with Dr. Freeman Lan in the design of the microfluidic and molecular workflow as well as Dr. Yu-yu Cheng and Dr. Ophelia Venturelli for providing strains. Integrated Training in Microbial Sciences (ITiMS) at the University of Michigan (Burroughs Wellcome) and National Science Foundation (Awards 2120909, 2426415) were critical in providing funding. This research was supported in part through computational resources and services provided by Advanced Research Computing at the University of Michigan, Ann Arbor.

## Data Availability Statement

Raw reads are available on NCBI Sequence Read Archive (SRA) (PRJNA1208117). Bioinformatic scripts for read processing and analysis are available on https://github.com/jamesyitan/Cocoa-seq-paper. Microscopy images, including fluorescence images of the bi-culture droplet communities and ddPCR droplets used for analyses are also available in relevant directories within the Github repository. Protocols.io link is available at https://www.protocols.io/view/combinatorial-co-culturing-and-amplicon-sequencing-n2bvj3nxblk5/v1.

