## Supplementary Information for "Highly multiplexed community profiling of sub-nanoliter droplet-based co-cultures"

**Extended Materials and Methods**

*Photolithography*

Microfluidic device designs (Fig. S1) (Supplementary Data 1 contains the raw AutoCAD designs) were made in AutoCAD, and transparency masks with these designs were purchased from Fine Line Imaging. SU-8 2050 (Kayaku Advanced Materials) was spincoated onto a 4-inch diameter silicon wafer to the appropriate thicknesses (40 µm and 80 µm) and baked pre-exposure at 65°C and 95°C for 6 and 18 minutes, respectively. Transparency masks with design features were aligned onto SU-8 coated wafers, put into contact with vacuum, and exposed to UV curing for 10 seconds. Wafers were then baked post-exposure at 65°C and 95°C for 4 and 12 minutes, respectively, and developed in SU-8 developer. Wafers were rinsed with SU-8 developer and isopropanol and blow dried. Device thicknesses were verified with a profilometer. The wafers were then hard baked at 175°C for 5 mins and rendered hydrophobic by vapor silanization with 2-3 droplets of tridecafluoro-1,1,2,2-tetrahydrooctyl-1-trichlorosilane in a vacuum desiccator for 1 hour before being baked again at 150°C for 10 minutes.

*Soft lithography*

The SU-8 master molds were taped to the bottoms of large petri dishes. PDMS elastomer base and curing agent mixed in a 9:1 ratio was poured on top of the master mold and degassed in a vacuum chamber to remove air bubbles. PDMS was cured for at least 8 hours at 65°C. After curing, the PDMS was peeled off and devices were cut out with a razor. Holes for device inlets and outlets were punched for each device using a 1.0 mm biopsy punch. After cleaning remaining debris on the devices with pressurized air and tape, devices were bonded onto glass microscopy slides (Fisher Scientific 12-518-101) with a plasma wand and baked at 80°C for 10 minutes. Device channels were rendered hydrophobic with Aquapel according to Mazutis et al. (1) and covered with scotch tape until use.

*Microbial cultures*

Benchmarking was performed with six microbial cultures: *E. coli* K12 BW25113 (metA::kan, pBbA2K-mCherry), *B. subtilis* 168 (trpC2, cat, amyE::Pveg-gfp-spec), *B. subtilis* 168 (amyE::Phyperspank-GFP(Sp), lacI), *Pseudomonas putida* KT2440, *B. thetaiotaomicron* VPI-5482, and *L. crispatus* ATCC 33820. Strains were maintained as glycerol stocks kept at -80°C. New cultures were inoculated from cryostocks into 5 mL of LB broth (Miller) in a 50 mL Falcon tube at 250 rpm overnight unless specified otherwise. *E. coli* was grown with 34 mg/L chloramphenicol at 37°C. Both *B. subtilis* strains were grown with 100 mg/L spectinomycin at 37°C. *P. putida* was grown at 30°C. *B. thetaiotaomicron* was grown overnight in RUM (modified YCFA) media (2) supplemented with 4 g/L fructose at 37°C anaerobically without shaking. *L. crispatus* was grown overnight in MRS media at 37°C anaerobically without shaking. Anaerobic growth was performed in an anaerobic chamber (Coy) with a 5% carbon dioxide, 2-4% hydrogen, and balance nitrogen atmosphere. 1 mL of each overnight culture was centrifuged in a 1.5 mL microcentrifuge tube at 4000xg for 5 minutes, the supernatant was removed, and the cell pellet was washed twice and resuspended in PBS. Each washed culture was appropriately diluted and counted with a C-Chip disposable haemocytomer (Fisher Scientific, #22-600-100) under a Nikon Ti-S microscope on the Ph2 phase contrast annulus with a 40X objective lens to quantify cell concentrations.

*E. coli*, *B. subtilis* 168 (trpC2, cat, amyE::Pveg-gfp-spec), *P. putida*, *B. thetaiotaomicron*, and *L. crispatus* were grown overnight individually and encapsulated in droplets (detailed in the next section) to test the cell lysis protocol for cell immobilized in agarose beads. The *E. coli* and *B. subtilis* 168 (amyE::Phyperspank-GFP(Sp), lacI) strains were used as a model obligately-syntrophic, two-species co-culture based off Hsu et al. (3). While the *E. coli* strain was the same as the one in Hsu et al., the *B. subtilis* is a modified strain with brighter GFP fluorescence. The *E. coli* and both *B. subtilis* strains were provided as a gift from the Venturelli Lab. When co-cultivated in droplets, the *E. coli*/*B. subtilis* co-culture was grown in M9, consisting of M9 salts (47.8 mM $Na_2HPO_4$, 22.0 mM $KH_2PO_4$, 8.55 mM NaCl, 9.35 mM $NH_4Cl$, 1 mM $MgSO_4$, 0.3 mM $CaCl_2$), micronutrients (2.91 nM $(NH_4)^2MoO_4$, 401.1 nM $H_3BO_3$, 30.3 nM $CoCl_2$, 9.61 nM $CuSO_4$, 51.4 nM $MnCl_2$, 6.1 nM $ZnSO_4$, 0.01 mM $FeSO_4$), thiamine HCl (3.32 µM) dextrose (D-glucose) at 5 g/L, and 2 mM IPTG. Three 4-member mock communities, comprised of *E. coli*, *B. subtilis* 168 (trpC2, cat, amyE::Pveg-gfp-spec), *P. putida*, and *B.*

*thetaiotaomicron* were mixed from individual, washed overnight cultures in designated ratios and co-encapsulated in droplets.

*Encapsulation of cells in agarose beads*

Droplet generation was performed in a modified large oven incubator (VWR Scientific 1535) to maintain a temperature between 37-40ºC to keep low-melting agarose suspensions fluid. To limit heat loss, we removed the oven incubator's outer door and replaced the inner glass door with an equivalent-sized, clear acrylic sheet (Optix, 30"x36"x.22") with holes cut by CNC milling (Digital Fabrication Lab, Taubman College of Architecture and Urban Planning, University of Michigan) to allow operators to adjust the microfluidic device and tubing as well as operate syringe pumps (Fig. S2).

3% agarose solution was prepared by melting low melting-point temperature (SeaPlaque GTG, Lonza) in PBS or medium in a 50 mL Falcon tube in a water bath at 70ºC, filtering the solution through a sterile 0.45 µm PDVF filter, and keeping the tube at 70ºC in the water bath until use. Right before encapsulation, the 3% agarose solution was mixed with 2X concentrated cell suspensions in a 1:1 volume ratio to obtain the appropriate cell suspensions in 1.5% agarose in 1X concentration of the respective media for droplet generation. Syringes with needles were preheated in the incubator or quickly with a heat gun on medium-low heat (Uline H-915). Around 600 µL of cell suspension were quickly withdrawn into a preheated sterile 1 mL syringe with a 23 G x 1" syringe needle (BD), air bubbles were removed, and the syringe was placed into the incubator oven to equilibrate with the interior temperature. PFTE tubing (Cole-Parmer, 0.022" ID x 0.042" OD) was used to connect the syringes to the inlets of the device as well as to collect generated droplet emulsions from the device outlet into a microcentrifuge tube. Two syringe pumps (Kent Scientific Genie Touch) in the incubator were used to flow the cell suspension and the oil phase comprised of HFE7500 Novec engineering oil with 2% fluorosurfactant (008-fluorosurfactant, RAN Biotechnologies, Inc.) into the droplet generation microfluidic device. The flow rate of both the oil phase and agarose suspension into the microfluidic device was 3 µL/minute. Approximately 200 µL of droplets were generated for each condition. If co-cultivation was performed, the collection tubes were transferred to a 37ºC incubator for 20-24 hours. Agarose in droplets was set by placing the collection tube on ice for 10-20 minutes.

In tests for lysis of cells immobilized in agarose beads, overnight cultures were individually suspended to a OD600 of 1.0 in 1X PBS and diluted 1:1 with molten agarose suspension (1X PBS, 17% Opti-Prep, 1% Pluronic F-68, 2% agarose) for droplet encapsulation and immediately set. Mock community cell suspensions were mixed from washed overnight cultures at twice the desired community compositions in 1x PBS and diluted 1:1 with molten agarose suspension (1X PBS, 30% Opti-Prep, 0.2% Pluronic F-68, and 3% agarose) for droplet encapsulation. The syntrophic model co-culture was mixed at twice the desired cell concentration in 1X M9 media and mixed 1:1 with molten agarose suspension (1x M9, 3% agarose). The syntrophic co-culture was co-cultivated after droplet generation before setting agarose.

After agarose was set, the oil phase below droplets in the collection tube was removed with a 200 µL gel loading tip. To break the droplets and collect agarose beads, we added 500 µL of PBS-wash buffer (1X PBS, 0.1% Triton X-100) and 500 µL of 20% (v/v) perfluorooctanol in HFE7500 oil to the droplet emulsion, vortexed well, centrifuged for 1 minute at 300xg, and removed the bottom oil phase. We then added 500 µL of 1% (v/v) Span-80 in hexane, vortexed well, and centrifuged for 1 minute at 300xg. The clear hexane layer on top was removed, and residual hexane was removed by adding 500 µL of PBS-wash, vortexing, centrifuging, and skimming off the milky hexane layer on top with a pipette tip. This was repeated 3 times until the supernatant was clear and the microgels formed a distinct pellet at the bottom. The suspension was mixed by pipetting and transferred to a clean 1.5 mL microcentrifuge tube. The tube was centrifuged for 1 min at 300xg, and supernatant was removed by pipettor, leaving the washed agarose beads.

*Microscope imaging to quantify co-culture composition*

To estimate *E. coli*/*B. subtilis* co-culture abundances, fluorescence images of hundreds of droplets were taken after co-cultivation. 5 µL of the droplet emulsion after co-cultivation and 5 µL of droplet oil were pipetted into a disposable haemocytomer (SKC, Inc. C-Chip™) and sealed with quick setting epoxy (Gorilla™ Epoxy, 4200102) to prevent evaporation. After waiting 30 minutes to 1 hour for the droplets to form a non-overlapping layer in the C-Chip, droplets were imaged with an Olympus DP71 fluorescence microscope with a color camera to capture both green and red fluorescence. In ImageJ (2.0.0-rc-69/1.52i), color channels were split to separate green and red images, and custom MATLAB (R2022a) scripts were used to detect droplets andnormalize fluorescence contributions from cells in droplets exaggerated from diffraction and quantify fluorescence of both strains in individual droplets. Droplets were detected from brightfield images which were processed in ImageJ to enhance the droplet boundary outline, which were detected as circles in MATLAB. Fluorescence normalization was performed by binarizing both red and green fluorescence images and converting larger red fluorescence blobs from the mCherry-*E.coli* strain which were present at lower cell density and more often on the peripherals of the droplets to the average size of the non-aggregate GFP-*B.subtilis* fluorescence blobs. Relative abundance for each strain per droplet was calculated for all imaged droplets by dividing total binarized fluorescence contribution from that strain in the droplet by the total binarized fluorescence contribute of both strains for the droplet.

All non-fluorescence microscopy was performed on a Nikon Eclipse Ti-S. Droplets were visualized in a similar manner with the C-Chip. For droplets with a diameter of 100μm or smaller, "Neubauer Improved" grid type was used, and for droplets with a diameter larger than 100μm, "Fuchs Rosenthal" grid type was used.

*Cell lysis of agarose beads*

Gels were washed with 1 mL 10 mM Tris-HCl three times; the resulting gel suspension was mixed with a 2X lysis buffer in a 1:1 ratio for a final concentration of 1 mM DTT, 1 mM Tris-HCl pH 8.0, 2.5 mM EDTA, 100 mM NaCl, 0.8% ready-lyse lysozyme (Lucigen, R1804M) and incubated in a 37°C shaker overnight. Lysis buffer was removed from gels after centrifugation, and gels were washed three times in 10 mM Tris-HCl pH 8.0. The resulting suspension was mixed with a 2X digestion buffer in a 1:1 ratio for a final concentration of 30 mM Tris-HCl pH 8.0, 10 mM EDTA, 0.8% Triton X-100 (v/v), 0.5% SDS, 1 µg/µL proteinase K (Lucigen, MPRK092) and incubated in a heat block for 30 minutes at 50°C. To deactivate proteinase K, gels were centrifuged; digestion buffer was removed; the remaining gels were washed in 10 mM Tris-HCl pH 8.0, 10 mM EDTA, 0.1% Tween-20 (v/v), 5 mM phenylmethylsulfonyl fluoride (PMSF) (Sigma-Aldrich 93482) and washed in 10 mM Tris-HCl pH 8.0, 10 mM EDTA, 0.1% Tween-20 (v/v) (TET buffer) five times; and stored at 4°C until use. Occasionally, a fine, white precipitate would form and settle below the agarose beads when the digestion buffer was added. The gels would be carefully removed by pipetting the microgel layer off into a separate microcentrifuge tube after centrifugation without removing precipitate.

To check that genomic DNA was immobilized in the microgels after cell lysis, 10 µL of packed microgels were incubated for 30 minutes in 1 mL of 10 mM Tris HCl with 1X SYBR Green stain in the dark. Afterwards, gels were pelleted by centrifugation, washed in TET buffer three times, and visualized in a disposable haemocytomer under fluorescence microscopy with a FITC filter.

*Barcoded hydrogel bead processing*

Barcode hydrogel beads were purchased as a unit of 1 million suspended in TET buffer (RAN Biotechnology, CustomSeqReady-1M, InDrop variation, ca. 147k). 16S 515f primer extension onto the barcode beads is based off the protocol from Zilonis et al. (4) for barcode extension on hydrogel beads. The purchased beads were centrifuged at 1000xg; washed three times in hydrogel bead wash buffer (5 mM Tris-HCl pH 8.0, 5 mM EDTA, 0.05% Tween 20); and incubated in 1X isothermal amplification buffer (NEB B0537S), Bst 2.0 DNA polymerase (350 U/mL) (NEB M0537S), dNTP mix (650 µM each), and 10 µM of the extension oligo comprised of the annealing region to the existing oligonucleotide and 16S 515f primer overhang (5'-TTACCGCGGCKGCTGRCACCTCCTGTCATCTCACTCCTG) for 1 hour at 60ºC protected from light. Beads were centrifuged, supernatant was removed, remaining beads were incubated in STOP-25 buffer (10 mM Tris-HCl pH 8.0, 25 mM EDTA, 0.1% Tween-20, 0.1 M KCl) at room temperature for 30 minutes and washed and incubated in STOP-10 buffer (10 mM Tris-HCl pH 8.0, 12.5 mM EDTA, 0.1% Tween-20, 0.1 M KCl) three times at room temperature. To remove any annealed oligo on the beads, beads were washed and incubated in denaturation solution (0.15 M NaOH, 0.5% Brij-35) three times and then washed with neutralization buffer (100 mM Tris-HCl pH 8.0, 10 mM EDTA, 0.1% Tween-20, 0.1 M NaCl) twice. Beads were washed three times with hydrogel bead wash buffer and three times with hybridization buffer (10 mM Tris-HCl pH 8.0, 0.1 mM EDTA, 0.1% Tween-20, 0.33 M KCl). Any unelongated oligonucleotides on the bead were removed by ExoI clean-up.

To do so, beads were incubated for 30 minutes with 20 µM of protective oligos complementary to the 16S 515f reverse region, then incubated in 1X ExoI buffer and ExoI (0.27 U/µL) (Thermo Fisher Scientific, EN0581) for two hours at room temperature to digest any single stranded oligos that were not hybridized to the protective oligo. To remove the protective oligos, beads were washed in STOP-25 buffer, incubated, washed in STOP-10 buffer three times, incubated and washed in denaturation buffer three times, washed in neutralization buffer twice, and washed in TET buffer three times before storage at 4ºC.

After the extension protocol, barcode beads were validated to ensure proper 16S forward primer addition on oligonucleotides with respective FAM oligo probes (IDT, 5-/56-FAM/TTACCGCGGCKGCTGRCAC for the 515f region and 5-/56-FAM/AGATCGGAAGAGCGTCGTGTAGGGAAAGAG for the PE1 region). Probes were annealed to the bead oligonucleotides at a concentration of 10 µM in 1 mL QC buffer (5 mM Tris-HCl pH 8.0, 5 mM EDTA, 0.05% Tween-20, 1 M KCl) in each tube, incubated for 20 minutes at room temperature in the dark, and washed with QC buffer three times to remove free FAM probes. Beads were concentrated with centrifugation and visualized on C-Chip cell count haemocytomers (SKC, Inc., DHCN015) with FITC filters.

*Unique molecular identifier (UMI) benchmarking*

The 16S standard is an oligonucleotide composed of a modified 16S V4 sequence from *Thermus thermophilus* ATCC 33923. The full sequence is as follows: 5-GTGCCAGCAGCCGCGGTAANNNNNNNGGCGCGAGCGTTACCCGGATTCACTGGGCGTAAAGGGCGTGTAGGCGGCCTGGGGCGTCCCATGTGAAAGACCACGGCTCAACCGTGGGGGAGCGTGGGATACGCTCAGGCTAGACGGTGGGAGAGGGTGGTGGAATTCCCGGAGTAGCGGTGAAATGCGCAGATACCGGGAGGAACGCCGATGGCGAAGGCAGCCACCTGGTCCACCCGTGACGCTGAGGCGCGAAAGCGTGGGGAGCAAACCGGATTAGATACCCGGGTAGTCC-3. The first 19 and last 20 nucleotides of the sequence are conserved to allow the standard 16S 515f and 16S 816r primers, respectively, to anneal, and the seven nucleotides directly after the 515f region are replaced with seven degenerate oligonucleotides for use as unique molecular identifiers. The 16S standard was purchased from GenScript as a single strand DNA (ssDNA) lyophilized stock with PAGE purification and verification for quality control. The ssDNA stock was resuspended in molecular-grade water and quantified with an in-house digital droplet polymerase chain reaction (ddPCR) to determine the concentration of functional 16S standard molecules. To do so, we diluted the stock $10^4$ to $10^9$ fold and made PCR reactions for each. The PCR reaction was composed of 0.6 µL of 816r 16S V4 reverse primer (10 µM), 0.6 µL of 515f 16S forward primer (10 µM), 5 µL of 5X SuperFi II Buffer, 0.6 µL of 10 mM dNTP (each) mixture, 0.6 µL of Platinum SuperFi II DNA polymerase, 0.375 µL of 20 mg/mL bovine serum albumin (BSA), 3 µL of 10% Pluronic F-68, 3 µL of 16S standard dilution, and 15.225 µL of molecular-grade water. This reaction volume was run through the droplet generation device with QX200 Droplet Generation Oil for EvaGreen (Biorad, cat # 1864005) to produce droplets with a diameter of approximately 40 µm with aqueous and oil flow rates of 10 and 60 µL/minutes, respectively. The droplets were collected in a 0.2 mL PCR tube for each dilution, excess oil was removed, and remaining droplets were covered with 50 µL of mineral oil. The droplets were thermocycled using the following program: 95ºC for 2 minutes, 98 for 30 seconds, 30 cycles of (98 for 10 seconds, 60 for 10 seconds, and 72 for 30 seconds) with a 2 ºC/s ramp rate between steps and no lid heating. Mineral oil was removed, and the oil phase was replaced with 10X SYBR with 2% surfactant in HFE7500 oil (0.2 µm filtered to remove excess SYBR in DMSO) and incubated for 20 minutes before two washes in fresh 2% surfactant in HFE7500 for 30 minutes. Droplets were viewed in a C-Chip under a FITC filter to identify the correct dilution that would provide a $\lambda$ of 0.1 template molecules/droplet. This condition was imaged, and images were processed with a custom MATLAB script to determine the concentration of functional 16S standard in the standard stock. The stock 16S standard concentration was approximately $2.91 \times 10^6$ molecules/nL.

*Droplet barcoding*

50 µL of 515f-extended barcode beads was washed in bead buffer (10 mM Tris HCl pH 8.0, 0.1% Tween 20, 50 mM KCl) in 0.5 mL PCR tubes by centrifugation at 1000xg for 1 minute three times, and a gel-loading pipette tip was used to remove as much liquid as possible after centrifugation at 3000xg for 2 minutes. gDNA-agarose beads were washed in 10 mM Tris-HCl pH 8.0 by centrifugation at 1000xg for 1 minute three times and as much liquid was removed with a gel-loading pipette tip. Packed agarose gels were diluted 1:1 by volume with 2X agarose bead suspension buffer (20% OptiPrep, 2% (w/v) Pluronic F-68, 0.6 mg/mL BSA, 20

mM Tris-HCl pH 8.0) for a 50% agarose bead suspension at 30 µL in a 0.5 mL PCR tube. 120 µL of PCR mastermix solution (0.3 mM of PE2-816r reverse primer, 1.5X SuperFi II buffer, 300 µM for each base of dNTPs, 3% Platinum SuperFi II DNA polymerase, 1.2% Pluronic F-68 (v/v), 0.36 mg/mL BSA) was prepared.

Nikon Ti-S inverted microscope was used to monitor droplet encapsulation of beads and microgels. A red band-pass filter (Midopt, BP635) was equipped into the condenser filter to prevent exposure of trace amount of UV-light from the microscope onto the UV-light sensitive barcode beads. Four syringe pumps (three GenieTouch syringe pumps from Kent Scientific and one KD Scientific) were used for microfluidic operation.

The barcoding device has four inlets for the PCR reagent carrier phase, the packed barcode beads, the agarose microgel suspension, and the oil, as well as one outlet for the droplets. Flow was controlled by a syringe pump holding an Air-Tite 1 mL plastic syringe filled with Biorad droplet generation oil and capped with a syringe needle. PFTE tubing was cut and a transfer syringe - an empty syringe with a needle - was inserted into one end of the tubing to manually draw an appropriate amount of each phase from a microcentrifuge tube into the other end of the PFTE tubing without drawing up air. Before withdrawing the barcode bead or agarose microgel suspension, the suspensions were pulse-vortexed 10 times to ensure homogenous distribution and break up clusters of gels or beads. The end of the tubing was removed from the transfer syringe and placed onto the end of the syringe and the needle filled completely with oil, without any air between the liquid interface and the needle syringe with oil. The other end of the PFTE tubing was attached to the inlet of the device. The syringe pump was run to slowly prime the liquid suspension to the inlet of the device at a steady flow of 10 µL/minutes. Packed barcode beads were loaded into the PFTE tubing and covered with a black tubing sheath (McMaster-Carr) to protect the barcode beads from light.

The flow rates for the syringe pumps were as follows: the oil flow was 5 µL/minutes, the barcode beads between 0.45-0.5 µL/minutes, the agarose suspension between 0.8-1 µL/minutes, and PCR reagent 3 µL/minutes.

For each condition, we collected droplets for 3 minutes from the outlet into 0.5 mL PCR tubes. For droplet thermocycling in 0.2 mL PCR tubes, each condition had 20 µL of Biorad oil and 30 µL of droplets from barcoding. Droplets were exposed to UV-light on ice under a 365-nm UV-light (Ted Pella Blak-Ray) for 10 minutes (with the lamp pre-heated for 10 minutes beforehand). 50 µL of mineral oil was placed on top of the droplets, and the reaction tubes were briefly centrifuged on a mini-centrifuge. Droplets were thermocycled using the following program: 95 ºC for 2 minutes; 98 ºC for 30 seconds; and 30 cycles of 98 ºC for 10 seconds, 60 ºC for 10 seconds, and 72 ºC for 30 seconds, with a 2 ºC/s ramp rate between all steps and no lid heating. Mineral oil was removed immediately after thermocycling finished and as much excess oil as possible was removed with a gel-loading pipette tip.

For PCR clean-up, 40 µL 1X ExoI buffer with 1 U/µL ExoI and 30 µL of perfluorooctanol was added for each condition and briefly centrifuged to merge droplets through incubation at 37 ºC for 30 minutes. Approximately 50 µL of aqueous phase was transferred without any oil to new 0.2 mL? PCR tubes for clean-up with AMPure XP beads and eluted in 20 µL of 10 mM Tris HCl 8.0.

*Library preparation and sequencing*

Library preparation was performed with PCR in a 50 µL reaction comprising of 2.5 µL of each library primer with appropriate indices (0.5 µM stock), 25 µL of 2X NEBNext Q5 HotStart HiFi PCR MasterMix, 0.125 µL of BSA (20 mg/mL), 9.875 µL of molecular grade water, and 10 µL of DNA clean-up elution. The thermocycler program was as follows: 98 ºC for 30 seconds; 5-7 cycles of 98 ºC for 10 seconds, 68 ºC for 20 seconds, and 65 ºC for 30 seconds; 65 ºC for 2 minutes; and a 12 ºC hold. The 490 bp PCR product was verified by running gel electrophoresis on a 1.5% agarose gel at 100 mV for 50 minutes, staining with ethidium bromide for __ minutes, and washing for 5 minutes in TAE buffer. Verified libraries were sent to the University of Michigan Advanced Genomics Core for quality control with Qubit and Agilent TapeStation and for sequencing with the Illumina NextSeq 2000 with P1 300 cycle chemistry. Paired end sequencing was performed with asymmetric read lengths (261/41 forward/reverse base pairs) according to the manufacturer's protocol (Illumina NextSeq1k2k). BCL Convert Conversion Software v4.0 (Illumina) was used to generate de-multiplexed Fastq files for each library.

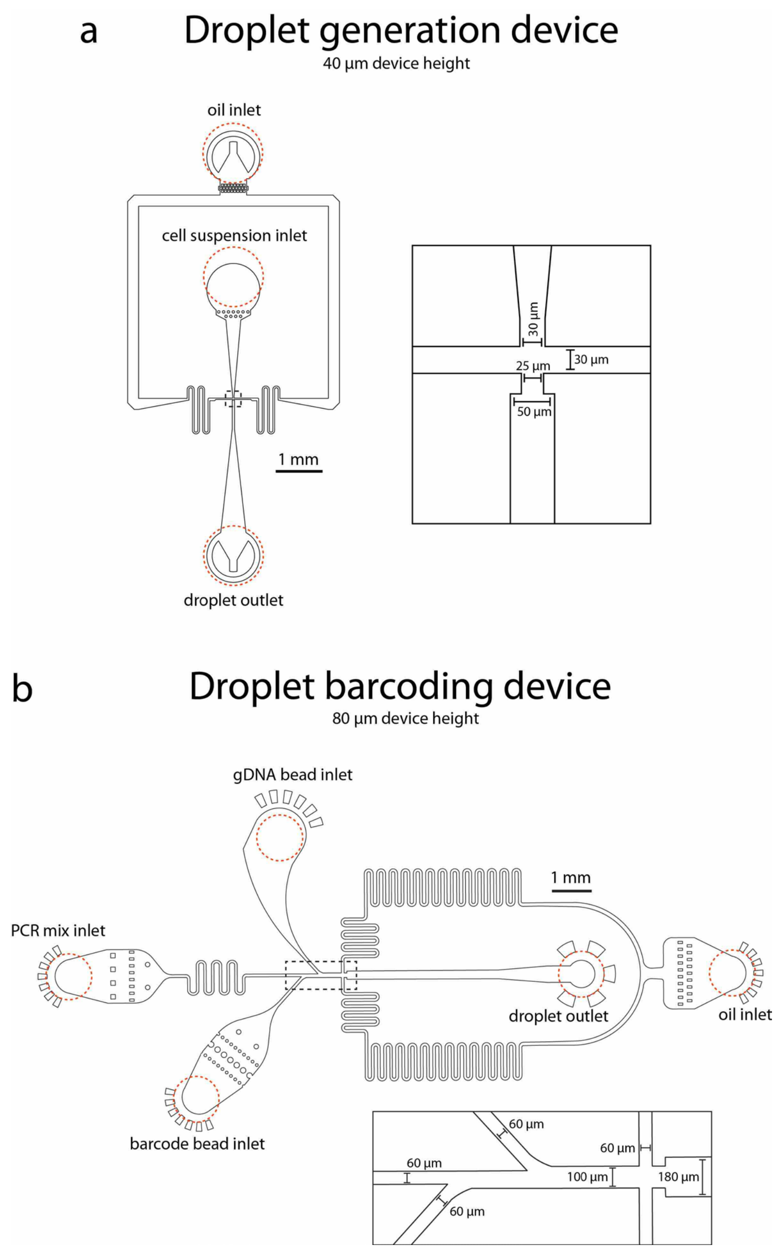


**Fig S1. Schematic of microfluidic PDMS devices used for droplet generation and droplet barcoding.** (a) The droplet generation device has an oil inlet, cell suspension inlet, flow-focusing channel intersection (dashed box), and droplet outlet. (b) The droplet barcoding device has an oil inlet, PCR mix inlet, gDNA bead inlet, barcode bead inlet, flow-focusing channel intersection (dashed box), and droplet outlet. Flow-focusing intersection sections in the dashed boxes are enhanced, and critical channel dimensions are specified. Device heights are also specified. The locations where inlet and outlet holes were punched for tubing are specified in dashed red circles. CAD designs are available in Supplementary Data 1.

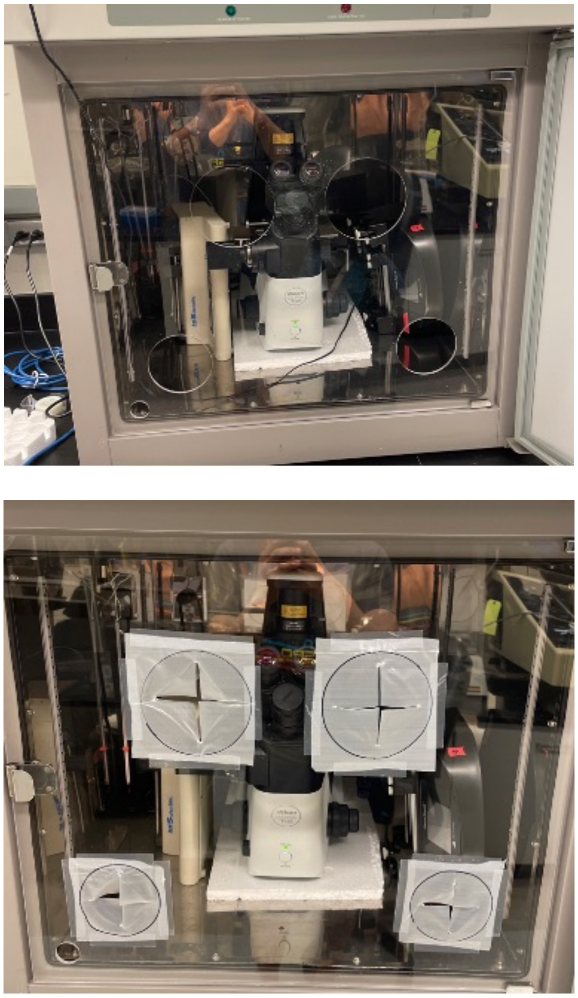

**Fig S2. Modified oven incubator for generating droplets with low-melting point agarose suspensions.** (Top) The original glass door of the oven incubator was replaced with a clear acrylic sheet of similar dimensions. The acrylic had holes cut out for entry of hands. (Bottom) Thin plastic sheets with slits were taped onto the holes to limit heat loss while hands were not inserted. The small uncovered hole at the bottom left allowed power cords to exit the incubator.

**Note S1.** Estimating number of co-cultures multiplexed for sequencing

The total number of communities multiplexed is determined from this equation:

$$N = \frac{0.74V\eta}{\left(\frac{4}{3}\pi\left(\frac{D}{2}\right)^3\right)}$$

Droplet number requires estimations of 3 parameters:

- Droplet size after multiplexing ($D$, diameter)
- Volume of droplets thermocycled ($V$)
- Efficiency of co-encapsulating gDNA beads with barcode beads ($\eta$)

Droplet size after multiplexing: This is determined by taking a small number of droplets (usually 1 µL) after multiplexing (before thermocycling) and viewing them in an improved Neubauer counting chamber (C-Chip™ Disposable Hemacytometer - Fisher Scientific 22-600-113) with 9 µL of droplet generation oil. The approximate diameter of droplets generated from Cocoa-seq is 150µm.

Volume of droplets thermocycled: The number of droplets processed is determined by the volume of droplets during droplet multiplexing that were thermocycled. Because the droplets float to the surface in a water-in-oil emulsion and have the potential to be well packed, we assume 74% packing efficiency (which is the highest theoretical packing efficiency of spheres in a three-dimensional space). Because droplets are manually pipetted into the PCR tube, the volume transferred is recorded and pipetted slowly to ensure droplets are well packed.

Co-encapsulation efficiency: This is determined by manually counting microscopy images in a hemocytometer after droplet multiplexing.

### Note S2. Microfluidic device design

The microfluidic design was based on the InDrops cell barcoding device (4) with the addition of a second bead inlet to co-encapsulate polyacrylamide barcode beads with agarose gDNA beads. Because the barcode bead has the same chemical composition as the InDrops bead, its inlet channel characteristics remain unchanged with the exception of “stopper columns,” which sequester polyacrylamide beads during initial flow until a sufficient packing factor is reached to ensure highly packed bead flow. However, the inlet for the agarose bead suspension was modified to accommodate the different mechanical properties of agarose beads. Agarose beads are much more compactable than polyacrylamide beads and, if compacted too much, can clump and flow irregularly. To prevent this, packed agarose beads are suspended in a 50% (v/v) suspension with a surfactant and BSA to prevent sticking. The inlet channel for agarose beads is also asymmetric and curved to prevent particle arching and clumping.

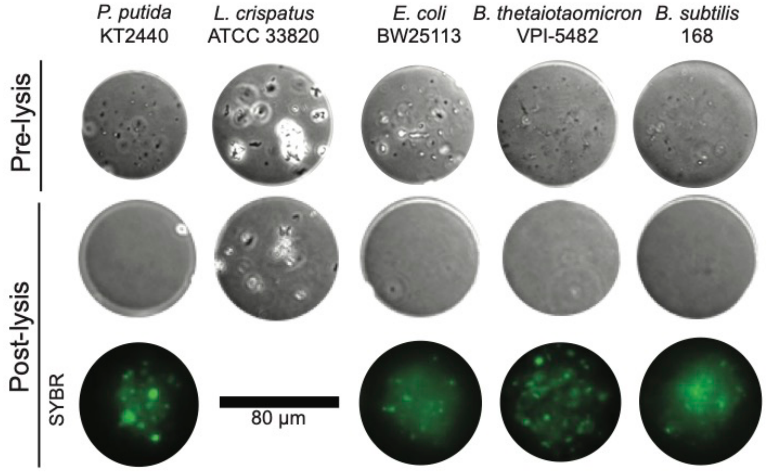


**Fig S3. Cell lysis of individual strains immobilized in agarose beads varies by cell type.** Beads pre-lysis are in the top row and beads that have undergone lysis are in the bottom two rows. Each visualized bead is representative of all beads observed for that condition. Due to difficulty imaging and processing the same bead in a large population, pre-lysis and post-lysis images are not of the same bead. However, SYBR images are of the same bead as the bright field image directly above it. No SYBR stain was performed for *L. crispatus* due to incomplete cell lysis.

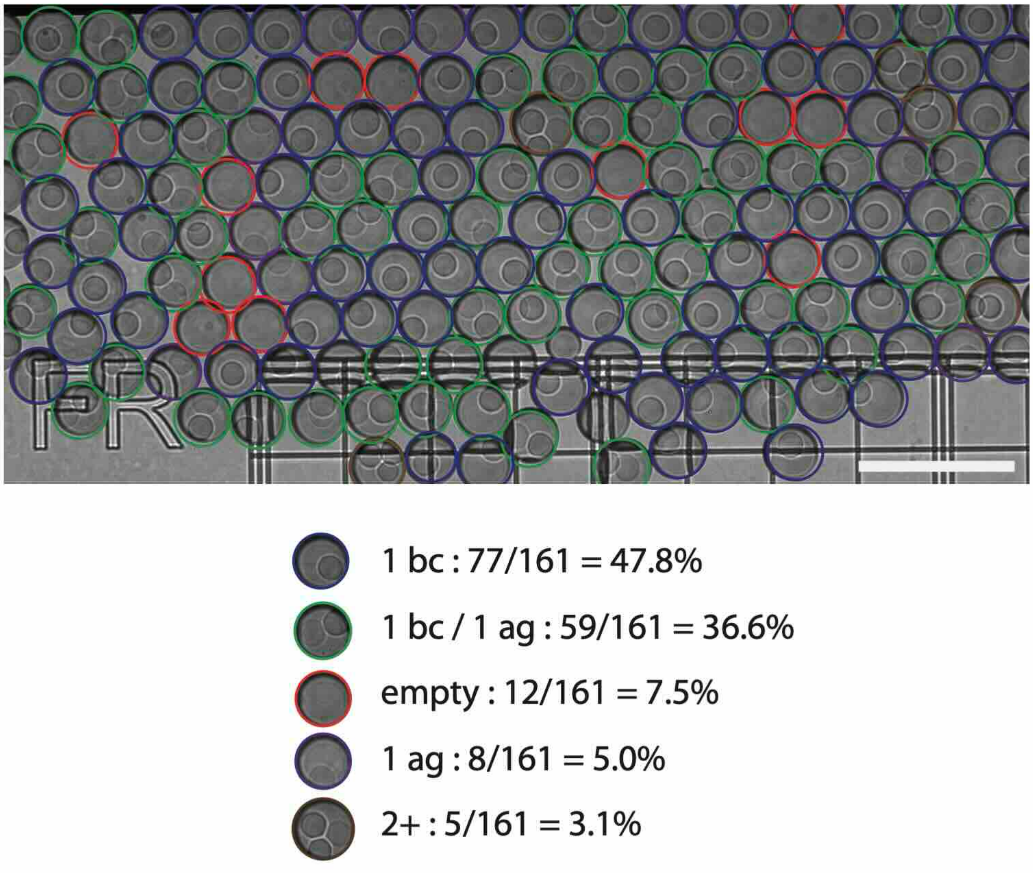


**Fig S4. Co-encapsulation efficiency of the droplet barcoding device.** A subset of the droplets generated from the droplet barcoding device was imaged and manually counted to determine co-encapsulation efficiency. Blue outlined droplets represent droplets with 1 barcode bead (1 bc), green outlined droplets represent desired droplets with exactly 1 barcode bead and 1 agarose bead (1 bc / 1 ag), red outlined droplets represent droplets with no bead (empty), purple outlined droplets represent droplets with just one agarose bead (1 ag), and brown outlined droplets represent droplets with 2 or more beads of the same type. Scale bar corresponds to 500 μm.

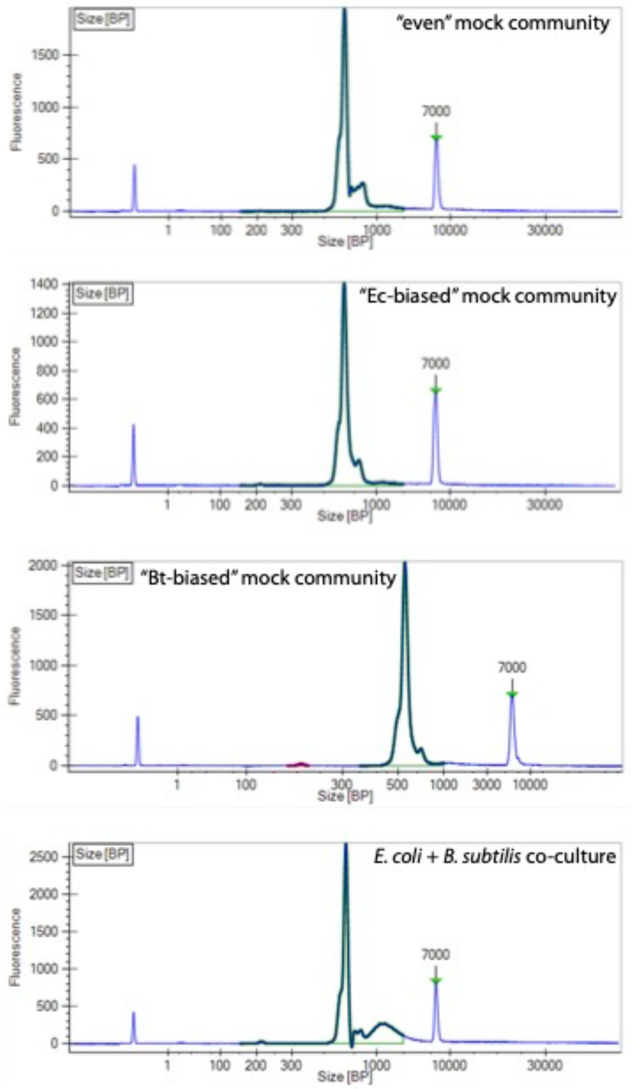


**Fig S5**. **Library quality confirmation by TapeStation after Cocoa-seq and Illumina library PCR**. The amplicon library peak is expected around 500 bp. The peak at 7000 bp is a gel migration marker.

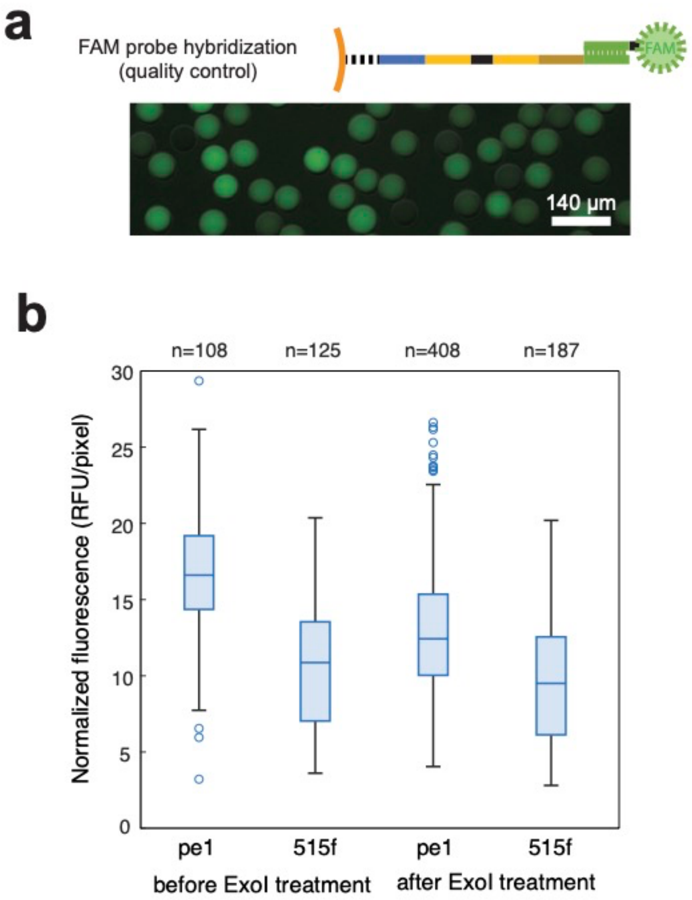


**Fig S6. Quality control of primer extension onto purchased BHBs.** (a) Hybridization of complementary fluorescent oligo probes confirms the presence of the 515f primer in elongated oligonucleotides on finished beads. (b) Quantification of oligo extension on barcode beads using fluorescent probes targeting the extended 515f region or the pe1 PCR handle site present in all oligos on the beads. Fluorescence values were normalized for each bead by dividing the total fluorescence by the size of the bead (total number of pixels). Comparison between the fluorescence of extended 515f regions and the fluorescence of pe1 region present on all beads (before ExoI treatment) reveals the efficiency of isothermal extension is similar to values observed in similar applications (4,5). Fluorescence values after ExoI treatment indicate that unextended oligos were mostly removed. The box plot indicates median, first, and third quartiles; whiskers indicate minimum and maximum, and points indicate outliers. The total number of beads quantified for each condition is labeled at the top of the graph.

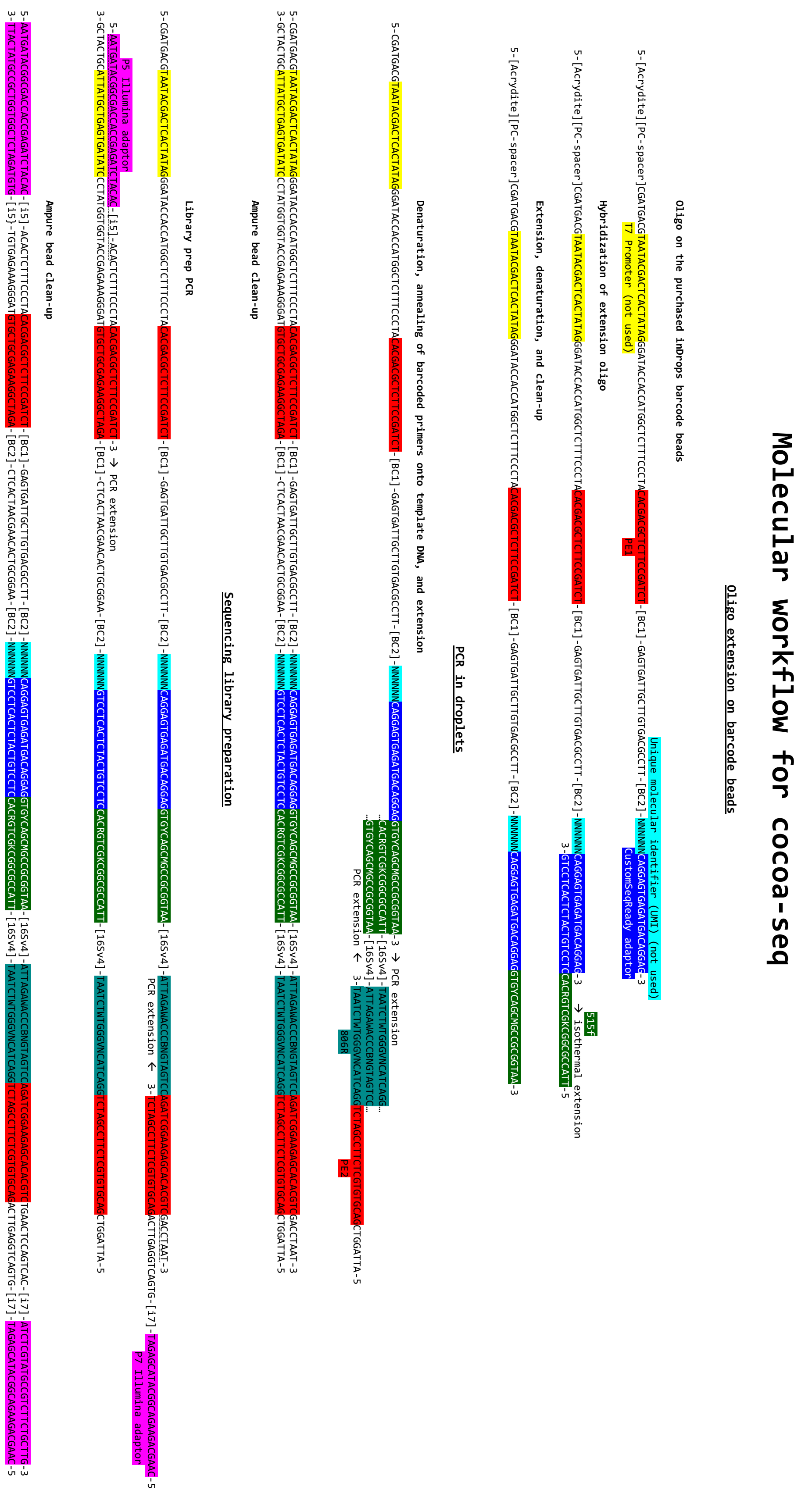


**Fig S7. Oligo design and molecular workflow for bead extension, droplet PCR (multiplexing), and Illumina library PCR.** Oligo regions are labelled with corresponded colored text. List of oligos used is available in the protocols.io entry.

| Sequencing library | Illumina indices | # reads | % of total sequencing run | % perfect w/ index | % one mismatch index reads |
|---|---|---|---|---|---|
| Even mock | CTCATCGA-TAGATCGC | 4483938 | 3.51 | 100 | 0 |
| Bt-biased mock | GGATCGTT-CTCTCTAT | 17271208 | 13.51 | 100 | 0 |
| Ec-biased mock | CCACGTTA-TATCCTCT | 21314159 | 16.67 | 100 | 0 |
| EcBs biculture | TCCTCCAT-CTCTCTAT | 40860298 | 31.96 | 100 | 0 |

**Table S1. Initial demultiplexing of sequencing libraries from Illumina indices**. All sequencing libraries were sequenced on the same run, as described as above. The remainder of reads which did not match the specified Illumina indices (34.35% of reads) was discarded.

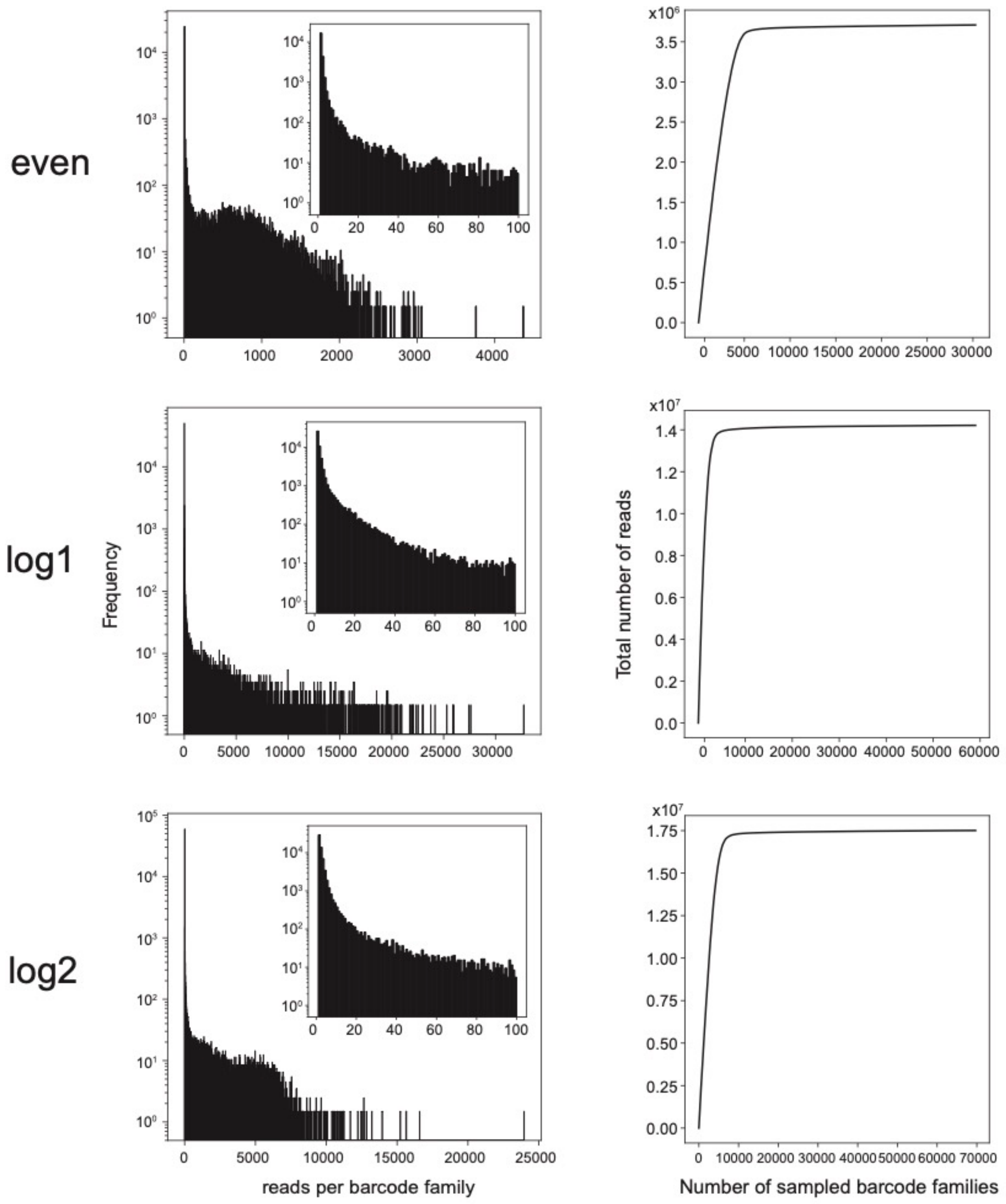


**Fig S8. Distribution of reads for mock community benchmarking libraries.** (Left) Histograms of the number of reads per barcode families with a bin size of 10. The subset histogram shows a more resolved view (bin size of 1) of the smallest barcode families. (Right) Rarefaction curve of the cumulative number of reads contributed from sampling barcoding families.

a

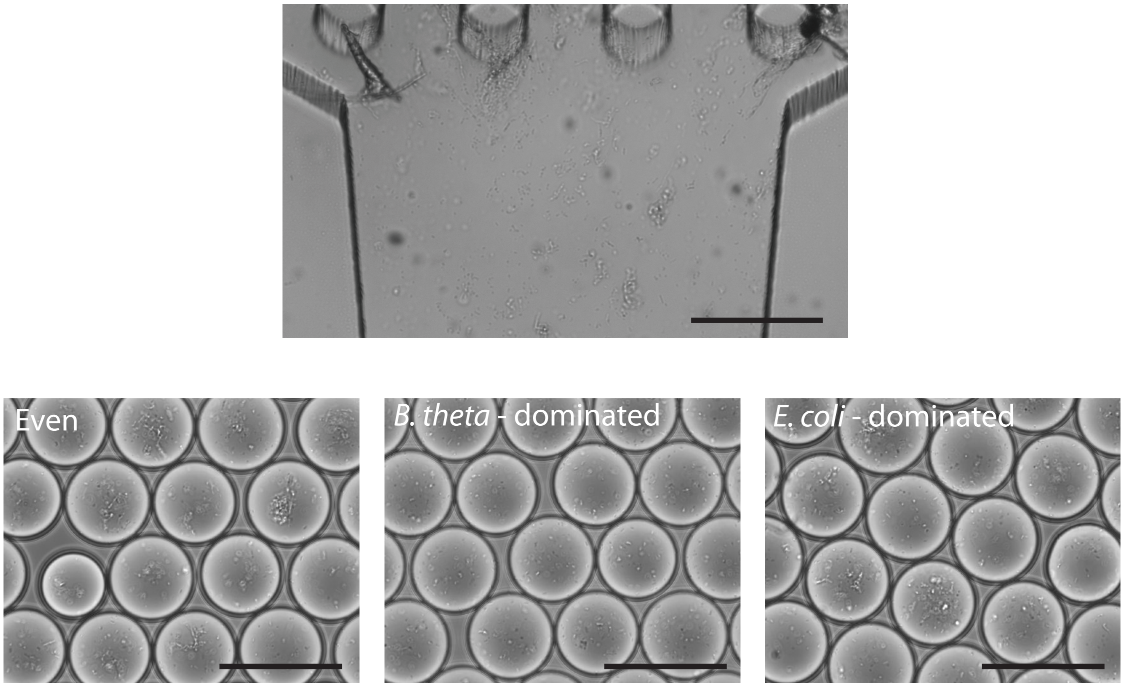


b

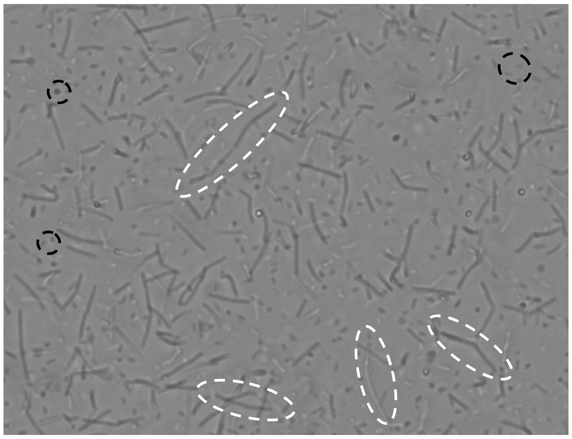

**Fig S9. Deviations from expected relative abundances in mock communities are caused by cell clumping and the disproportionally large size of *B. subtilis* cells.** (a) Cell clumping was observed in the microfluidic channels before droplet encapsulation (top) and inside droplets after encapsulation (bottom). The *B. thetaiotaomicron*-dominated community has the least amount of clumping, but extensive clumping is present in the “even” and *E. coli*-dominated mock communities. Scale bar is 100μm. (b) Microscopy image of the “even” mock community. Cells of this specific *B. subtilis* strain grow as long rods and are distinguishable from the rest of the cells in the community. Some *B. subtilis* are identified (white dashed circles) and smaller cells from the other members in the community are identified (black dashed circles).

| | *Bacillus subtilis* | | *Bacteroides thetaiotaomicron* | | *Escherichia coli* | | *Pseudomonas putida* | |
|---|---|---|---|---|---|---|---|---|
| | Cocoa-seq | In-silico | Cocoa-seq | In-silico | Cocoa-seq | In-silico | Cocoa-seq | In-silico |
| Even mock | 0.53±0.12 | 0.34±0.05 | 0.14±0.09 | 0.17±0.03 | 0.20±0.09 | 0.24±0.04 | 0.13±0.08 | 0.24±0.04 |
| Bt-biased mock | 0.05±0.13 | 0.02±0.02 | 0.79±0.26 | 0.86±0.04 | 0.004±0.04 | 0.003±0.006 | 0.15±0.24 | 0.12±0.04 |
| Ec-biased mock | 0.34±0.09 | 0.12±0.04 | 0.002±0.007 | 0.001±0.003 | 0.65±0.09 | 0.87±0.04 | 0.006±0.02 | 0.008±0.009 |

**Table S2. Average relative abundances (± standard deviation) of each species in various droplet mock communities in Fig. 5b.**

| | *Bacillus subtilis* | | *Bacteroides thetaiotaomicron* | | *Escherichia coli* | | *Pseudomonas putida* | |
|---|---|---|---|---|---|---|---|---|
| | Cocoa-seq | In-silico | Cocoa-seq | In-silico | Cocoa-seq | In-silico | Cocoa-seq | In-silico |
| Even mock | 1.00 | 1.00 | 0.967 | 1.00 | 0.998 | 1.00 | 0.981 | 1.00 |
| Bt-biased mock | 0.734 | 0.559 | 1.00 | 1.00 | 0.213 | 0.186 | 0.910 | 1.00 |
| Ec-biased mock | 0.996 | 0.999 | 0.228 | 0.196 | 1.00 | 1.00 | 0.731 | 0.542 |

**Table S3. Prevalence of each species in various droplet mock communities in Fig. 5b.**

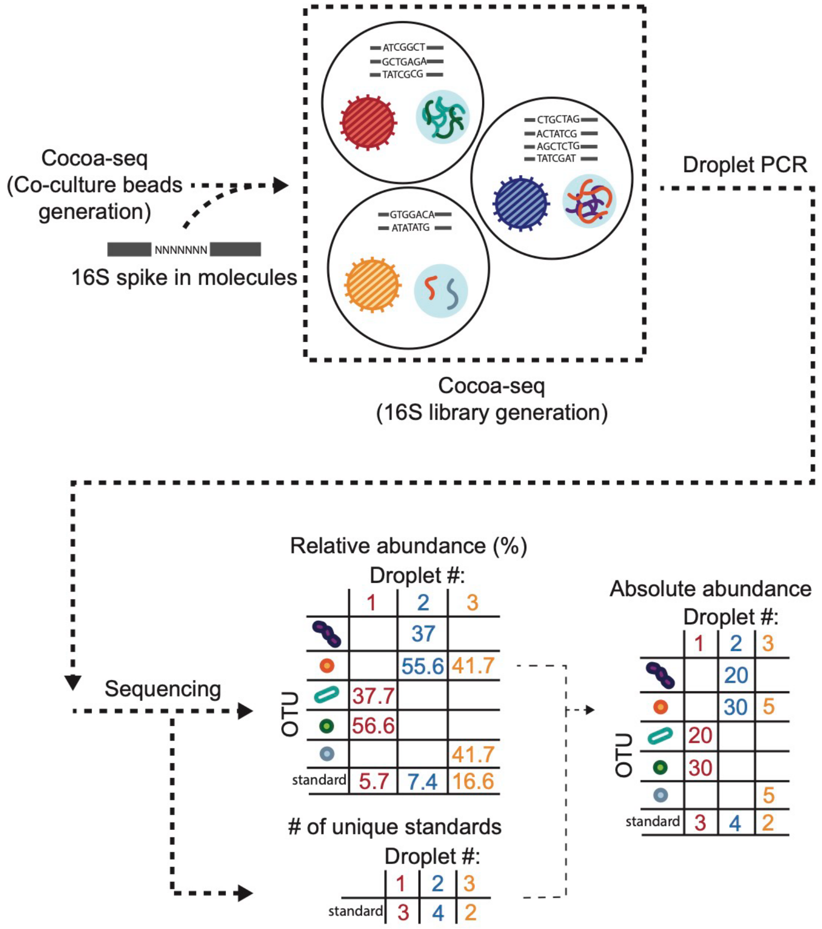


**Fig S10. Proposed workflow for determining absolute abundance of droplet communities using spike-in molecular standards.** Molecular standards are synthetic oligonucleotides designed with a specific 16S V4 sequence with a 7-mer degeneracy and are included in Cocoa-seq during the secondary droplet co-encapsulation for droplet library multiplexing. After Cocoa-seq, these standards are identified for each individual library and used to calculate absolute abundance from relative abundance.

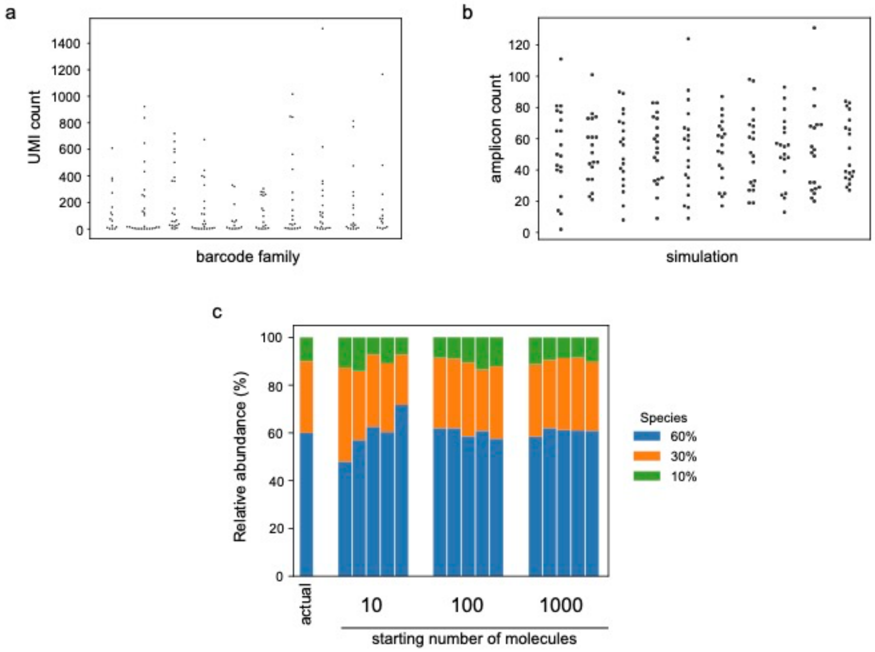


**Fig S11. Observed and modelled stochastic PCR amplification of individual initial templates.** (a) Observed distributions of the count of each UMI after PCR amplification in different droplets (barcode family) during the Cocoa-seq workflow. Initial droplet encapsulation of molecules was at a λ of 50 molecules/droplet. UMIs present less than 2 times in each droplet are not included. (b) *In silico* distributions of counts of discrete molecules after simulated PCR from 10 simulations. All simulations started with 20 molecules that underwent 20 cycles of amplification with a subsampling of 1000 molecules. (c) *In silico* amplifications of an uneven community comprised of three members (at 60-30-10% relative abundance) with different initial numbers of starting molecules (10, 100, or 1000). All amplifications were subsampled to 1000 molecules. Each condition had 5 replicates.
